# Dynamic spatiotemporal coordination of neural stem cell fate decisions through local feedback in the adult vertebrate brain

**DOI:** 10.1101/2020.07.15.205021

**Authors:** Nicolas Dray, Laure Mancini, Udi Binshtok, Felix Cheysson, Willy Supatto, Pierre Mahou, Sébastien Bedu, Sara Ortica, Monika Krecsmarik, Sébastien Herbert, Jean-Baptiste Masson, Jean-Yves Tinevez, Gabriel Lang, Emmanuel Beaurepaire, David Sprinzak, Laure Bally-Cuif

## Abstract

Neural stem cell (NSC) populations persist in the adult vertebrate brain over a life time, and their homeostasis is controlled at the population level. The nature and properties of these coordination mechanisms remain unknown. Here we combine dynamic imaging of entire NSC populations in their *in vivo* niche over weeks, pharmacological manipulations, mathematical modeling and spatial statistics, and demonstrate that NSCs use spatiotemporally resolved local feedbacks to coordinate their decision to divide. These involve a Notch-mediated inhibition from transient neural progenitors, and a dispersion effect from dividing NSCs themselves, exerted with a delay of 9-12 days. Simulations from a stochastic NSC lattice model capturing these interactions demonstrate that they are linked by lineage progression and control the spatiotemporal distribution of output neurons. These results highlight how local and temporally delayed interactions occurring between brain germinal cells generate self-propagating dynamics that maintain NSC population homeostasis with specific spatiotemporal correlations.

## INTRODUCTION

The maintenance of organ physiology over a lifetime in adult vertebrates is permitted by the activity of resident stem cells (SCs). Adult SCs locally generate differentiated progeny cells directed towards functional plasticity, cell replacement or organ growth. Adult SCs also self-renew, to ensure their own maintenance. Because differentiation and self-renewal occur concomitantly, the perdurance of SC populations (hereafter referred to as homeostasis) is a dynamic process. It remains however poorly understood how adult SCs achieve this long-term, dynamic and spatiotemporal equilibrium.

In a number of adult epithelial systems, SC clonal behavior appears compatible with stochastic decisions of gain or loss taken among equipotent SCs, indicating that SC numbers are maintained at the population level (Greulich and Simons, 2016; Klein and Simons, 2011; Rulands and Simons, 2016; Simons and Clevers, 2011). In addition, another important aspect of SC pools homeostasis is the control of SC fate choices in space. Indeed, the positioning of SC recruitment events will impact the location of progeny cells, hence organ growth and function. Such population-based homeostasis suggests the existence of feedback control mechanisms coordinating stemness-related fate choices over time (Lander, 2009; Rue and Martinez Arias, 2015) and in space. These feedbacks remain however largely unexplored at the molecular level in adult SC systems. From the spatial perspective, a prevalent example of coordination was illustrated in mouse interfollicular epithelial SCs, where the delamination of differentiating SCs triggers division of neighboring SCs in response to mechanical stretching (Mesa et al., 2018). It remains however unknown whether these findings can be generalized to other SC systems, in particular systems with much slower turn-over than epithelia and where SCs may not be prone to immediate division.

Along these lines, a particularly relevant case is the vertebrate adult brain, where neural stem cells (NSCs) are mostly quiescent, with in average one recruitment event every few weeks or months (Basak et al., 2018; Than-Trong et al., 2020; Urbán et al., 2019). This is added to a generally deep location inside the brain, making NSC pools very difficult to study dynamically in situ (Pilz et al., 2018). In niches of the adult pallium in zebrafish and mouse (sub-ependymal zone of the lateral ventricle -SEZ-, sub-granular zone of the dendate gyrus -SGZ-), NSCs are apico-basally polarized astroglial cells, arranged in neuroepithelium-like assemblies from which their neuronal progeny emigrates (Gonçalves et al., 2016; Obernier et al., 2018; Than-Trong and Bally-Cuif, 2015). NSCs can stochastically choose distinct fates, whose equilibrium is maintained through population asymmetry (Basak et al., 2018; Than-Trong et al., 2020). However, we lack an integrated spatiotemporal understanding of NSC population behavior, and of how their homeostasis can be dynamically maintained in time and space at long term.

To address this issue, we made use of the everted organization of the adult zebrafish pallium, where NSCs are organized as a dorsally located ventricular monolayer (Adolf et al., 2006; Grandel et al., 2006). Using transparent adult mutants and transgenic backgrounds reporting NSCs or cell states, it is possible to image NSCs in their niche, and reconstruct their behavior over weeks (Barbosa et al., 2015; Dray et al., 2015). We applied it here to reconstruct the tracks of all NSCs within entire portions of the dorsal pallium and study, at the population level, the spatiotemporal regulation of the most upstream NSC fate decision: activation. Activation is the transition from the quiescent to the proliferating state, and signals NSC recruitment. In the zebrafish adult pallium at any given time, NSC activation events are broadly distributed across the entire NSC pool (Dray et al., 2015). Using long-term intravital imaging of the dorso-medial pallium (Dm), we reveal that NSC activation events are not randomly positioned but respond to cell-cell inhibitory cues that operate over space and time within the NSC pool. From quantitative experimental parameters, we develop a mathematical modeling platform that faithfully recapitulates NSC population behavior, and demonstrate that the spatiotemporal dynamics and propagation pattern of NSC recruitment events is an emergent property of lineage cues. These analyses provide the first quantitative understanding of spatiotemporal homeostasis in adult vertebrate NSC pools.

## RESULTS

### NSC activation events are randomly positioned relative to each other across the NSC population at any given time

Adult pallial NSCs in zebrafish -and rodents-are radial glial cells expressing Glial Fibrillary Acidic Protein (Gfap) (Ganz et al., 2010; März et al., 2010). Whole-mount immunostainings for GFP in *Tg(gfap:gfp)* transgenic adults (Bernardos and Raymond, 2006) (3 months post-fertilization, mpf), together with the Proliferating Cell Nuclear Antigen (PCNA), reveal the distribution of proliferating cells at large scale within the pallial NSC pool (Figures 1A and B), and three essential ventricular cell states/types (from thereon referred to as “states”): quiescent NSCs (qNSCs) in the G0 state (GFAP+;PCNA-), activated NSCs (aNSCs) in G1-S-G2-M (GFAP+;PCNA+) and activated neural progenitors (aNPs) (GFAP-;PCNA+). These cells are lineage-related (Figure 1C). qNSCs activate periodically and can return to quiescence, while aNPs are generated by symmetric or asymmetric neurogenic divisions of aNSCs, and will delaminate from the germinal sheet to generate neurons (Rothenaigner et al., 2011; Than-Trong et al., 2020). At any time, a triple labeling for Gfap, PCNA and the general progenitor marker Sox2 shows that aNSCs represent 5.2 % of the entire progenitor population in Dm (NSCs + aNPs), against 82.5 % qNSCs and 12.3 % aNPs (Figures 1D, S1A-D and Table S1).

**Figure 1.**
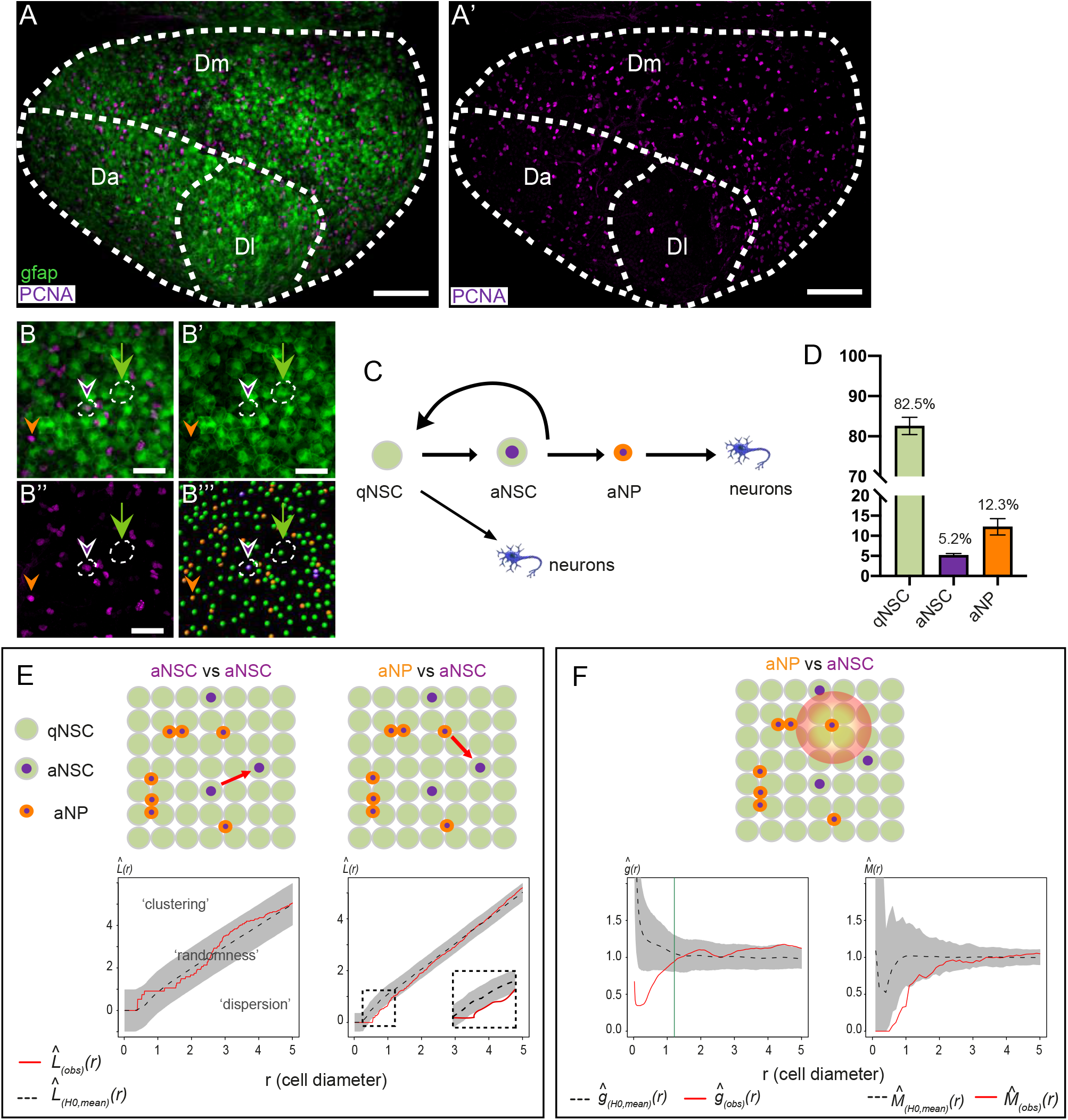
Inhibitory interactions between progenitor cells bias the distribution of NSC activation events. **(A, A’)** Confocal whole mount view of the pallial germinal layer in a 3mpf *Tg(gfap:GFP)* fish stained for GFP (green, NSCs) and PCNA (magenta, proliferating cells). Anterior left, pallial subdivisions (Dl: lateral, Dm: medial and Da: anterior) indicated by dotted lines. **(B-B’’’)** Close-up in Dm showing progenitor cell states: quiescent NSCs (qNSCs; GFP+ only; green arrow), activated NSCs (aNSCs; GFP+,PCNA+; magenta arrowhead), proliferating neural progenitors (aNPs; PCNA+ only; orange arrowhead). **B**: Merge; **B’,B”**: individual channels; **B’’’**: segmented image (green: qNSCs, magenta: aNSCs, orange: aNPs). **(C)** Main pallial NSC lineage (arrows: lineage transitions). **(D)** Proportions of qNSCs, aNSCs and aNPs relative to each other in Dm. **(E)** Besag’s *L*-function assessing spatial correlations for the same fish between aNSCs (left) and aNSCs and aNPs (right). *L_(obs)(r)_* (red line): experimental value, *L_(H0,mean)_(r)* (black dotted line): mean under the Random Labelling null hypothesis (the state of any cell is independent of other cells and of its position), *r*: mean cell diameter in this brain, grey regions: 95% confidence envelopes. High magnification shows significant dispersion. **(F)** Range and strength of this interaction determined with functions *g* (left) and *M* (right). aNSCs are at least four times less frequent within 1-cell diameter of an aNP than in the whole Dm. Green bars: 95^th^ centile of the distance to the furthest direct neighbor of aNPs. Scale bars: A-A’=100μm, B-B’’’=30μm.

Detection of every individual cell based on the position of its nucleus, located apically, generates an array of spatial coordinates that captures the position and the state of each cell at the studied time point (Figures 1B-B’’’). Full scale maps plotting local cell densities, cell states and their ratio across entire pallial germinal sheets highlighted strong differences between pallial subdivisions but similarities between hemispheres in individual fish (Figures S1E-E’’’), as well as between fish, arguing for the reliability of our whole-mount cell detection approach. In the remainder, we analyzed one hemisphere per animal.

To determine whether NSC activation events respond to specific spatial rules, as opposed to uniquely resulting from cell autonomous decisions, we next set out to quantitatively describe their distribution within the NSC population. First, because the PCNA protein is detected throughout the cell cycle in this system, including during shortly post-M phase, we filtered PCNA-positive signals to distinguish novel activation events (preceding cytokinesis), which are of interest here, from post-cytokinesis doublets of sister cells (Figure S2, Methods section “Filtering PCNA signals”). In the following, we refer to “NSC activation event” as the PCNA^+^ phase preceding division (Figure S2H). We next used a spatial point pattern analysis (Baddeley et al., 2016) to test for spatial interactions between NSC activation events within the pallial germinal sheet (Figure S3A, Methods section “Spatial statistics”). First, we tested if aNSCs were randomly distributed or depended on the position of other aNSCs at the same time point. To detect spatial correlations, we used Besag’s *L* functions (Besag, 1977), a variance-stabilized version of Ripley’s *K*-function (Ripley, 1977) with an isotropic edge correction. If a spatial interaction was detected, we turned to two second-order functions, *g* and *M* (Figure S3A). The pair correlation function *g* is a rescaled derivative of the *K*-function, necessary to determine the interaction range given that the *L*-function is cumulative. The *M*-function (Marcon and Puech, 2010; Marcon, Puech, and Traissac, 2012) is a recently developed extension of the *K*-function that is adjusted for cell frequencies across the entire Dm domain, offering easily interpretable values to quantify interactions. Specifically, the locations of aNSCs were compared with a random distribution under the Random Labelling null hypothesis, obtained by permuting the same number of aNSCs among the experimental number of NSCs (qNSCs + aNSCs) for each fish (Figure 1E). In each case, the 95% confidence envelopes delimit a pattern within which the distribution of aNSCs is considered random, as opposed to clustered or dispersed (Figure S3A). In all cases analyzed, the distribution of aNSCs relative to each other was found positioned within the random interval (Figures 1E and S3B-C). Thus, at any given time, NSC activation events are spatially independent from each other across the pallial germinal sheet.

### NSC activation events are locally inhibited by further committed progenitors along the neurogenesis lineage

Because aNPs could also provide local positional cues, we next assessed the position of aNSCs relative to aNPs, using Besag’s *L*-function. For each animal, the experimental distribution of aNSCs relative to aNPs was challenged against simulations where the same number of aNSCs is randomly permuted among all NSCs, while aNPs are maintained at their endogenous position. In this case, we observed a deviation from the random pattern, aNSCs being further away from aNPs than expected by chance (Figures 1E and S3D).

This dispersion may reflect inhibitory interactions between the two cell types. To further characterize the range and strength of this effect, we used Ripley’s *g* and *M* functions, respectively (Marcon and Puech, 2010; Marcon et al., 2012). Ripley’s *g* function revealed a statistically significant shift of aNSC positioning relative to aNPs for radiuses within 1 cell diameter (Figure 1F), indicative of short-range interaction. It falls within random chance boundaries for distances beyond the furthest direct neighbor. Within a one-cell diameter distance, Ripley’s *M* function revealed an interaction strength of 0.25-0.5 between aNSCs and aNPs, showing that aNSCs are found next to aNPs two to four times less frequently than expected by chance (Figure 1F). All four animals studied qualitatively and quantitatively showed the same interactions (Figure S3D). Together, these results indicate that, at any given time, NSC activation events tend to avoid a territory located in the immediate neighborhood of aNPs.

### The local inhibition of NSC activation by aNPs is Notch signaling-dependent

In the adult zebrafish pallium and mouse SGZ, Notch3 signaling promotes NSC quiescence (Alunni et al., 2013; Kawai et al., 2017). We previously reported that aNPs express the Notch ligand DeltaA (Chapouton et al., 2010). On whole-mount preparations, we now quantified that 99% of aNPs express the *deltaA:egfp* transgene at any time (Figures 2A and 2B). We thus tested whether aNPs could locally inhibit NSC activation via Notch-mediated lateral inhibition.

To conditionally decrease Notch signaling in the adult pallial germinal zone, we subjected 3mpf adults to a 24-hour treatment with the gamma-secretase inhibitor LY411575 (hereafter referred to as LY) (Alunni et al., 2013). This short treatment minimally affects cell fate/state: it only partially reactivates NSCs (4.1% aNSCs among NSCs+NPs in DMSO-treated controls, 4.8% in LY-treated animals) and is too short for most aNPs to differentiate (9.1% aNPs in DMSO-treated controls, 10.8% in LY-treated animals) (Figure 2C and Table S1). To also control that LY treatment did not affect aNP fate/state and distribution, we compared the position of aNPs relative to each other in LY-versus DMSO-treated animals using Besag’s *L* and Ripley’s *g* and *M* functions (Figures S3E and S4D). In all cases, aNPs appeared clustered at short range, with 2 to 5 times more aNPs within a 1-cell diameter range than expected by chance. This result reflects rapid amplifying divisions by a fraction of aNPs, generating small clusters prior to differentiation (Figure S2B). Neither this pattern nor its range and strength were affected by LY.

**Figure 2.**
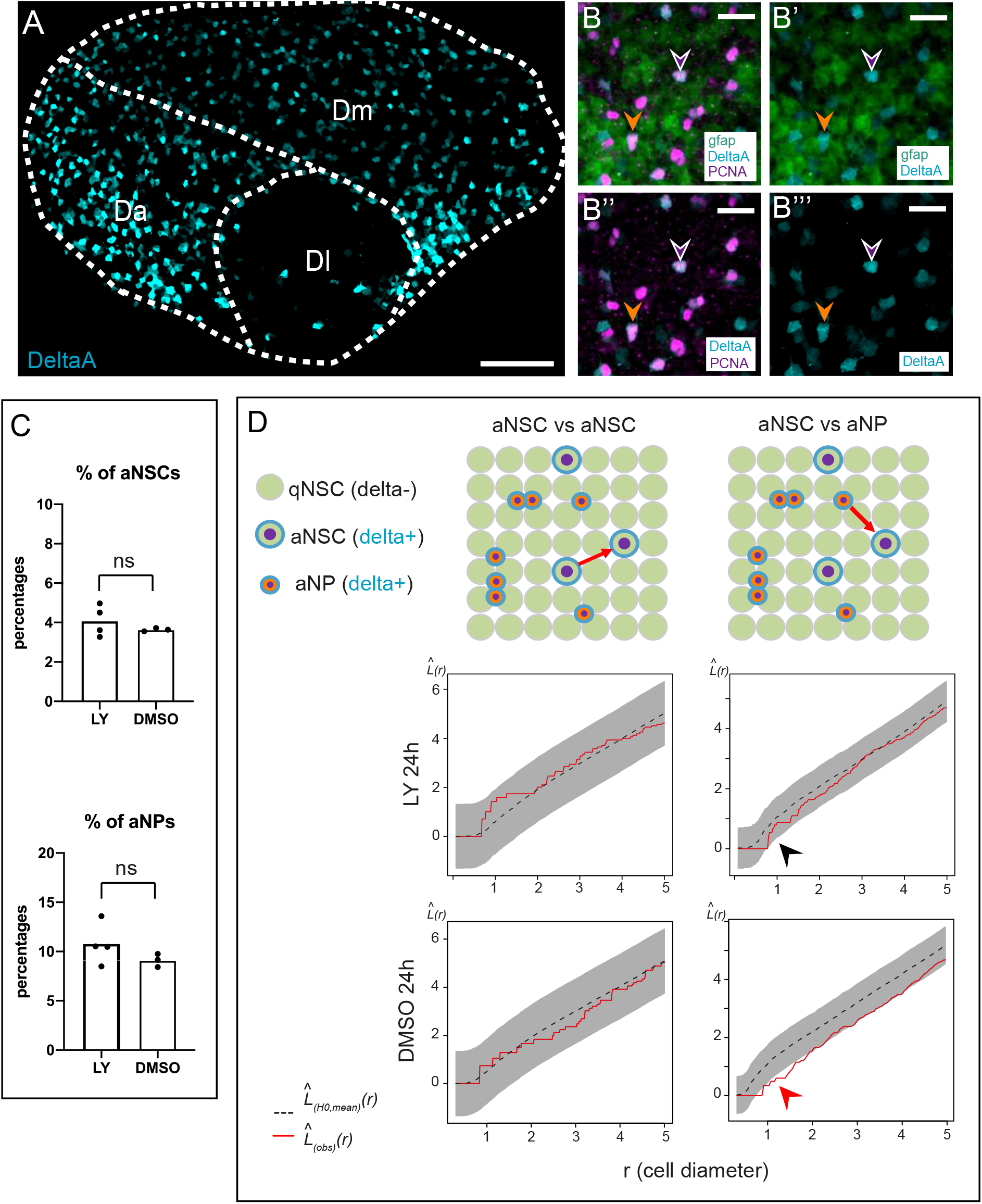
The inhibition of NSC activation exerted by aNPs is Notch signaling-dependent. **(A)** Confocal whole mount view of the pallial germinal layer in a 3mpf *Tg(gfap:tdTomato);Tg(delta:GFP)* fish stained for tdTomato (NSCs, green), GFP *(deltaA* expression, cyan) and PCNA (magenta) (DeltaA channel only). Anterior left. **(B-B’’’)** Close-ups in Dm, all channels. Magenta arrowhead: aNSC, and orange arrowhead: aNP, expressing *deltaA*. **(C)** Percentages of aNSCs and aNPs among qNSCs+aNSCs+aNPs upon 24h LY treatment (DMSO: control). **(D)** Spatial distribution (Besag’s *L*-function) of aNSCs relative to each other (left) and to aNPs (right) upon 24h LY treatment (DMSO: control). Red arrow: aNP inhibitory effect, black arrow: disappearance upon LY treatment. Scale bars: A=40μm, B-B’’’=20μm.

We next used this experimental scheme to assess whether the inhibition of NSC activation by aNPs was Notch-dependent. Besag’s *L*, Ripley’s *g* and Marcon’s *M* functions confirmed the short-range inhibition exerted by aNPs in all DMSO-treated controls (Figure 2D bottom right, Figure S4C). In striking contrast, this inhibition was abolished in all cases upon LY treatment (Figure 2D top right, Figure S4C). Thus, the inhibitory effect of aNPs on NSCs activation is, directly or indirectly, mediated by Notch signaling. Notch blockade was, in contrast, without effect on the random pattern of aNSCs relative to each other (Figure S4B).

Together, these results indicate that, at any given time, the spatial positioning of NSC activation events across the adult pallial germinal sheet responds to short-range Notch-dependent inhibitory cues exerted by aNPs, which decrease the probability for NSCs to activate in their immediate vicinity.

### NSC activation events are spatially controlled over time by preceding spatial activation patterns

Because homeostasis of SC ensembles is a dynamic process, we next addressed whether the spatial pattern of NSC activation events incorporates temporal information. Previous results indicate that the majority of aNSCs return to quiescence after division (Than-Trong et al., 2020). Thus, aNSCs observed at a few days interval reflect the successive activation of different qNSCs (Alunni et al., 2013). Using intravital imaging (Dray et al., 2015), we recorded NSCs in 3mpf transparent double mutant *casper* adults, double transgenic for *gfap:mTomato* (highlighting NSCs) and *mcm5:egfp* (highlighting activation events), in their endogenous pallial niche over 23 days at 3-4 days interval (Figures 3A and 3B). Next, in each animal (n=3), the position of every individual NSC of a complete ensemble of 370-500 neighboring NSCs was detected, and movies were assembled to reconstruct the behavior of every cell in the context of its neighbors (Figure 3C,C’ and Video S1). Hence, lineage trees (referred to “tracks”) could be produced for each NSC (Figures 3C’, S5A and S5B, Table S2). These tracks were filtered to focus on activation events preceding the first division (hereafter referred to as “NSC activation events”, like in the static analysis above) (Figure S5C, Table S2). As expected given the high proportion of quiescent NSCs and the slow dynamic of the system, most cell tracks were silent during the 3-4 weeks of recording: in average, 117 over 1203 tracks captured (9.7% +/− 0.8) led to at least a division event (Figure S5B). Most importantly, because we recorded every NSC of the population, these events were spatiotemporally resolved relative to neighboring cells (Figure 3C,C’), providing the first complete 4D dataset of NSC population behavior *in situ*. It is to note that, due to relatively weak *mcm5:egfp* staining, we could not efficiently resolve aNPs, which were therefore not considered in the dynamic analysis.

**Figure 3.**
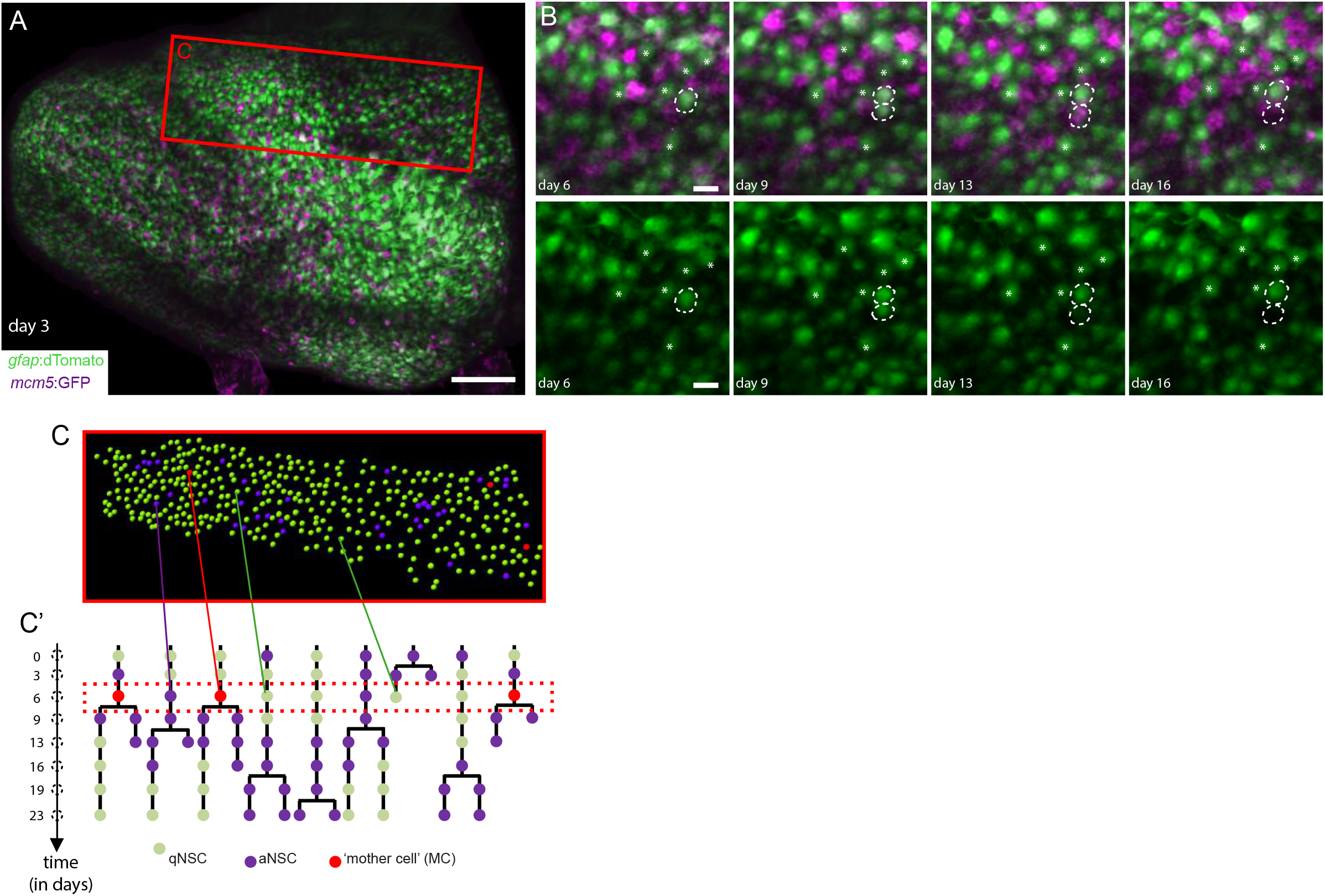
Intravital imaging resolves adult NSC lineage trees in time and space. **(A)** Whole pallial hemisphere imaged intravitally in a 3mpf *casper;Tg(gfap:dTomato);Tg(mcm5:GFP)* fish (individual named Mimi) (anterior left, image taken at d3 from a 35-day session of recordings every 3-4 days). Colors of the live reporters were fit to Figs. 1 and 2 (green: NSCs, magenta: proliferating cells). **(B)** Close-ups from the same movie showing an asymmetric NSC division (dotted circles) between d3 and d9: one daughter differentiates over the next 7 days (bottom dotted circle, loss of the *gfap:dTomato* signal). White asterisks: random qNSCs close to the division, used for alignment. **(C-C’)** Segmentation and NSC tracking over 23 days in Dm in Mimi. **C**: Segmentation of about 390 cells per time point (area boxed in A). **C’**: Example of dividing tracks, with cell states (color-coded) and the spatial position of each tree.

To validate the reliability of this imaging, cell detection and tracking approach, we first tested whether it could recapitulate the results obtained on the static pattern of aNSC placement. We applied a point pattern analysis at every individual time point to address the relative position of activation events within the whole NSC population. As previously concluded, these events were found randomly positioned relative to each other (Figures 4A and B). This analysis further revealed that the NSC activation phase can span several consecutive time points prior to division. We could indeed fit the NSC activated phase with an exponential decreasing function with a decay rate of 0.217 ± 0.019 day^−1^, corresponding to a mean aNSC half-life of 3.2 days (Figure S6B). Thus, in the following, we focused on the more temporally restricted event of cell division proper, reflecting the NSC recruitment event affecting cell fate. We refer to this event as the “mother cell” state (MC), defined as the imaged time point immediately preceding cell division) (Figures 3C,C’, 4 and S5A, red cells). MC events also appeared randomly positioned relative to each other at any given time (Figure 4C).

**Figure 4.**
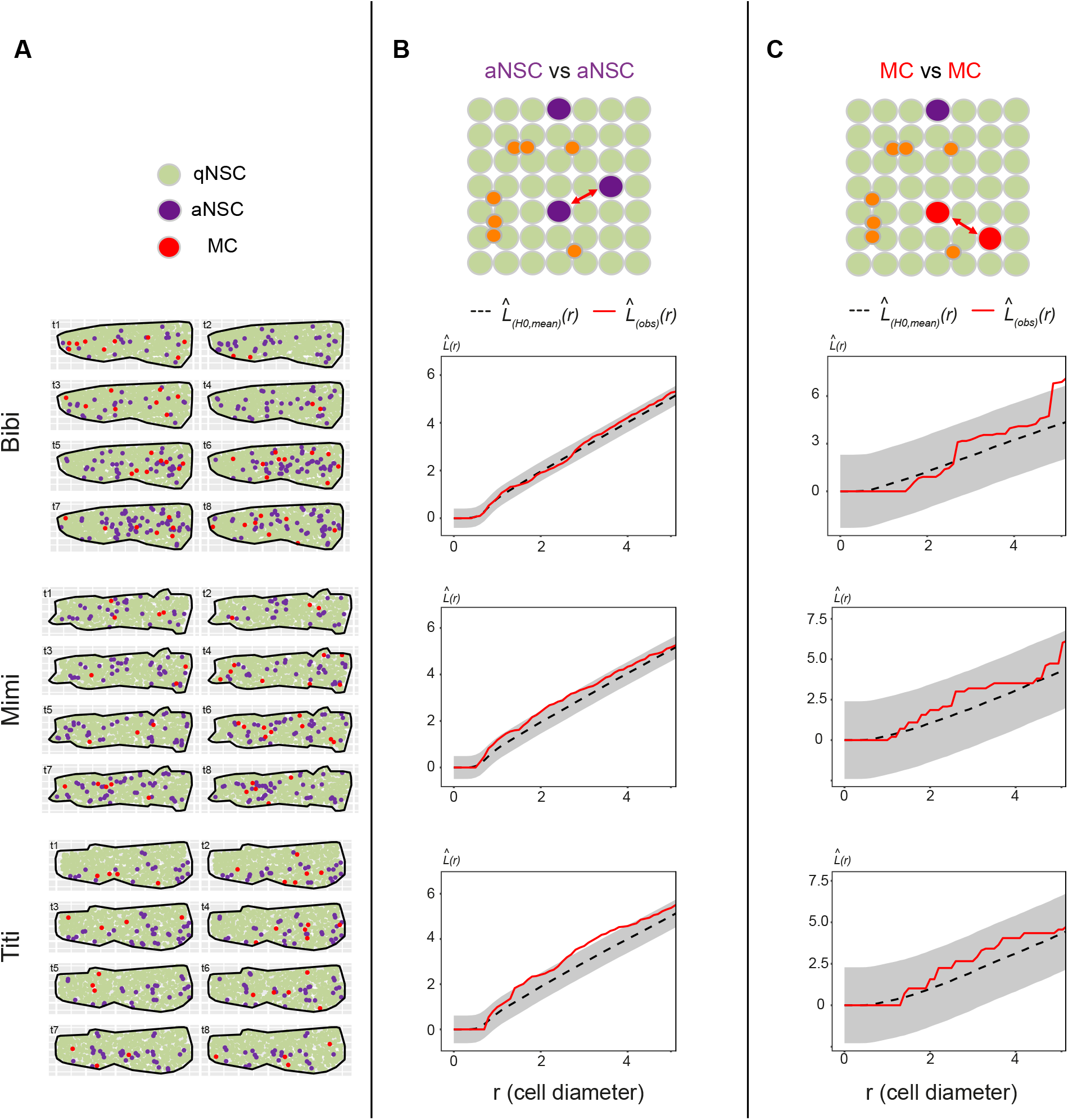
Validation of the dynamic cell detection and tracking method in a static spatial analysis of NSC activation. **(A)** Dm surfaces segmented for the three fish analyzed (Bibi, Mimi and Titi) at all time points, cell states color-coded. **(B-C)** *L*-functions respectively comparing the positions of aNSC and MCs with each other for each fish. *L_(obs)_(r)* (red lines): experimental values, *L*-functions for all time points pooled using a weighted average; *L_(H0,mean)_(r)* (black dotted lines): means under the Random Labelling null hypothesis; *r*: mean NSC diameter for each Dm surface; grey regions: 95% confidence envelopes.

To probe for the existence of temporal correlations, we next addressed whether MC positioning was influenced by the distribution of MCs at previous time points. In a point pattern analysis across time, we measured the position of MCs relative to the position of MCs at earlier imaging days at all possible time intervals (Δt1: 1 imaging time point = 3 days, Δt2: 2 imaging time points = 6 days etc), and confronted this to a scenario where the distribution of MCs at time t is random within all NSCs of the germinal sheet (Figure 5A). Strikingly, this analysis revealed a trend dispersion from the random distribution at Δt4 (interval of 4 imaging time points, i.e. 9-12 days) (Figure 5B). This trend is visible in all fish, is not observed at any other time interval, and reaches significance when pooling the 3 fish analyzed (>1200 cells per time point) (Figure 5B). Besag’s *L* integrated squared deviation and Ripley’s *g* function further indicate an effect exerted at a 1-to 2-cell diameter range, in all fish (Figure 5C). Thus, MCs at any given time are found further away than expected by chance from the position occupied by MCs 9-12 days earlier. Thus, the positioning of NSC division events incorporates spatiotemporal information with a delay, being less frequent within a radius of 1-to 2-cell diameter from the position occupied by dividing NSCs 9 to 12 days before.

**Figure 5.**
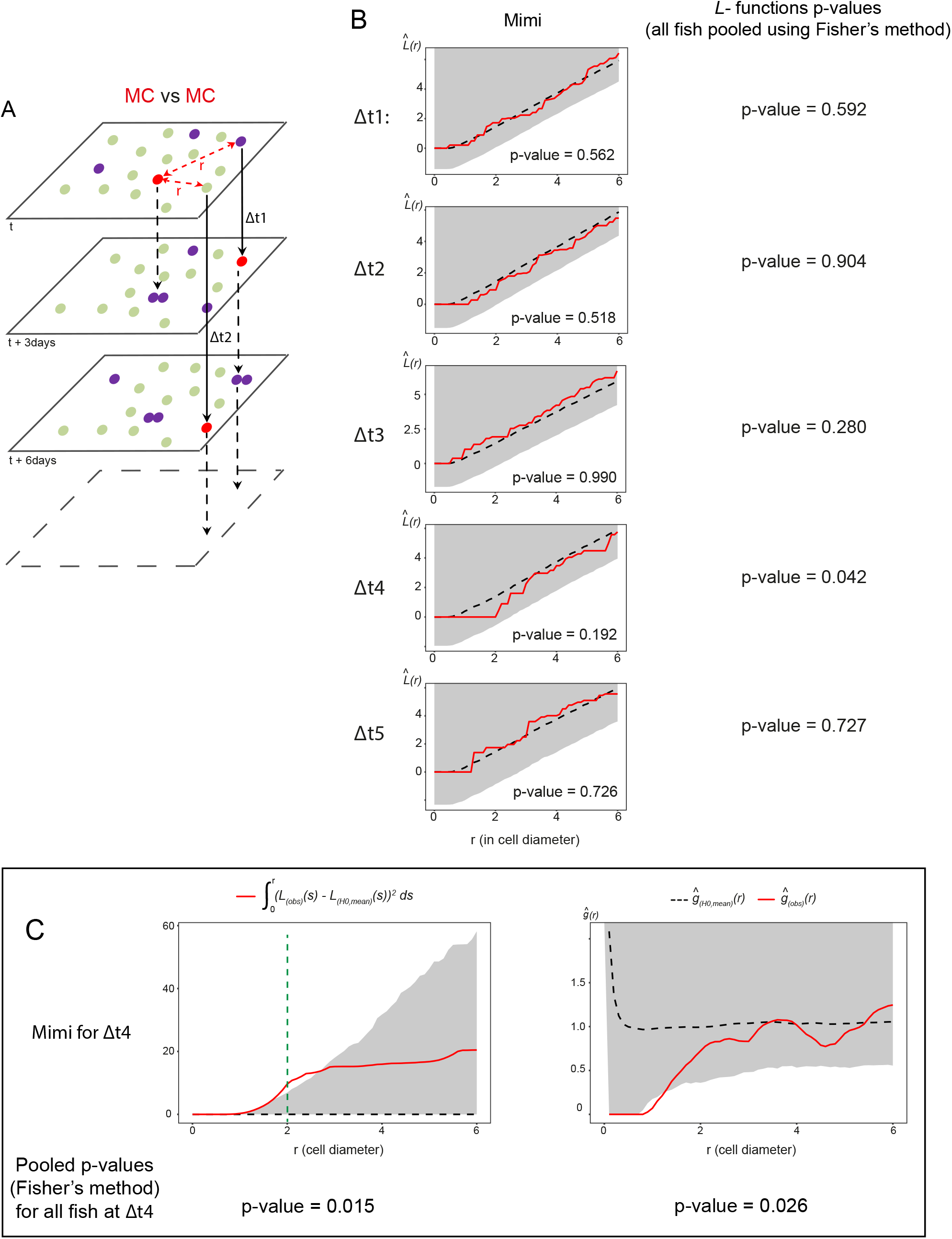
NSC division events are spatiotemporally coordinated. **(A)** MC positions at one time point t are compared using point pattern statistics with MC positions other time points after fixed intervals (t+Δt), for each possible Δt (3, 6, 9, and 15 days). **(B)** Left: Besag’s *L*-functions comparing MCs with each other for all Δt intervals in Mimi. *L_(obs)_(r)* (red lines): experimental values; *L_(H0,mean)_(r)* (black dotted lines): means under the Random Labelling null hypothesis; *r*: mean NSC cell diameter for each Dm surface: grey regions: one-sided 95% confidence envelopes. Each fish shows a trend dispersion at Δt4. Right: Combined *p*-values for Mimi, Bibi and Titi (Fisher’s method) (significant at Δt4). **(C)** Left: integrated square of the negative deviation between *L_(obs)_(r)* and *L_(H0,mean)_(r)* (surface under *L_(H0,mean)_(r)*) in Mimi at Δt4. Right: Ripley’s *g*-function indicating an effect at the nearest neighbor’s range. Top: illustrations for Mimi, bottom *p*-values, at two-cell diameter (green dotted line), obtained by combining the results from the three fish using Fisher’s method.

### A mathematical model of NSC behavior reveals the influence of population feedbacks in the spatiotemporal control of NSC activation and division events

The results above show that, within the adult pallial NSC pool, de novo NSC activation events follow spatiotemporal correlation rules encoded by the NSC/NP population: they are disfavored in the immediate vicinity of aNPs at all times, and, with a temporal delay of 9-12 days, in territories neighboring the position of previous NSC division events.

To gain insight into whether these events were linked, and whether they reflected *bona fide* instructive cues as opposed to emergent cues from the NSC ensemble, we developed a stochastic spatiotemporal model of NSC behavior (‘NSC lattice model’) (Figure S7A). The experimental input (Figure S6) includes the proportions of qNSCs, aNSCs, and aNPs at steady state, the duration of the NSC activation phase, division frequencies, and spatial information on the number qNSCs in contact with aNPs. Our previous clonal analyses demonstrated long-term homeostasis of NSC numbers within a tracked NSC population, with stable NSC proliferation rate and fates until approx. 15-18mpf (Than-Trong et al., 2020). Thus, the first layer of the model is a steady state mean field analysis of dynamic rate equations describing the transitions between the different progenitor cell states and differentiated neurons (Figures 6A and S7A, left), identifying transition rates (*γ* parameters in Figure 6A, Tables S3 and S4) based on quantitative experimental data. These rates are used as input for the second layer of the model (Figure S7A, right), which uses spatial stochastic simulations to obtain insight into the spatiotemporal correlations between the different cell states. The model uses a 2D vertex modeling platform to simulate transition between cell states, cell divisions, and cellular differentiation in a disordered cell lattice (Figure 6B). Cellular morphologies are determined by minimizing mechanical energy following these events (Figures S7B-G).

**Figure 6.**
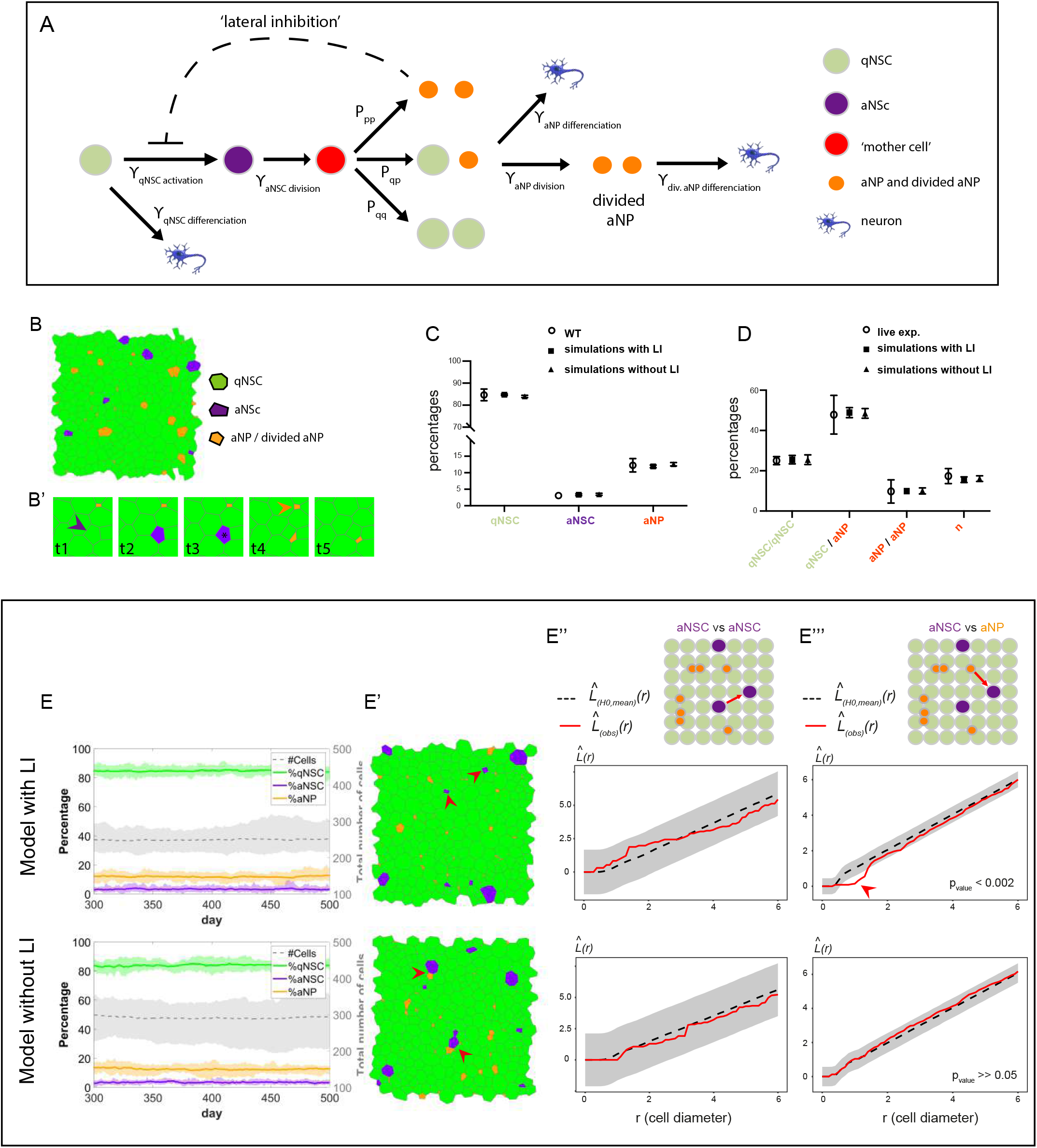
A ‘NSC lattice model’ captures NSC population dynamics. **(A)** Model lineage flowchart. γ: transition rates; P_qq_, P_qp_ and P_pp_: probabilities for symmetric and asymmetric aNSC divisions; dashed line: lateral inhibition (LI) of aNPs on NSC activation. **(B)** Snapshot of the lattice and **(B’)** examples of activation (purple arrowhead), division (black star) and differentiation (orange arrowhead) (Video S2). **(C)** Average cell proportions and **(D)** percentages of cell fates following cell division in simulations vs in vivo. **(E-E’’’)** Dynamics and spatial correlations from the NSC lattice model with (top) and without (bottom) aNP-driven LI. **E**: Stable cell numbers and proportions over time, matching experimental data. Bold line: mean; shade: STD; n=18 simulations, results of 1 simulation are shown. **E’**: Snapshots of the lattice at one time step from a simulation of 500 time steps. The simulation without LI shows many more cases where aNSCs neighbor aNPs (read arrowheads). **E’’,E’’’**: Besag’s *L* function of aNSCs relative to each other (**E’’**) and relative to aNPs (**E’’**). Red arrowhead to the dispersion of aNSCs from aNPs at a range of 1-2-cell diameter with LI.

Next, we introduced in the model the aNP-mediated inhibition of NSC activation (referred to as lateral inhibition -LI-) by suppressing the transition from qNSC to aNSC when the qNSC is in a direct contact with an aNP (dashed line in Figure 6A). To be able to compare between models with and without LI to assess spatial distributions, we also encoded the average effect of LI on the qNSC transition rates: we introduced a constant factor, *I*, that multiplied the qNSC rate, and compensated for the overall mean field effect of LI. This factor is estimated from the average fraction of qNSCs that are not in contact with any aNP (Figure S6F).

Long-term simulations over hundreds of days in the models with and without LI showed that stability is reached within tens of days when starting from random initial distributions. Simulations analyzed between days 300 and 500 showed maintenance of steady state proportions of cell states and total cell numbers (Figure 6E), as well as average cell sizes and number of neighbors (not shown) (Video S2). The qNSCs, aNCs, and aNPs fractions at steady state, and the percentages of cell fates following cell divisions or direct differentiation, match experimentally measured fractions (Figures 6C, D and E). These simulations also recapitulated the spatial bias in aNSC distribution relative to aNPs at any time, with similar range and strength as observed in vivo, in the model run with but not without LI (Figures 6E, E’ and E’’’). All other static spatial correlations uncovered in vivo were also observed: aNSCs appear randomly positioned relative to each other at any time, and aNPs appear locally clustered due to their rapid divisions (Figure 6E” and not shown). Thus, the LI interaction exerted by aNPs on NSC activation encoded in the model is sufficient to faithfully reproduce the in vivo relative distributions of progenitor cell states at any time.

Next, we used the NSC lattice model to shed light on the origin and determinant parameters of other spatiotemporal interactions detected within the adult pallial NSC population. Along these lines, the delayed effect of MCs on future MC events could reflect an active action of MCs, or emerge from the spatiotemporal characteristics encoded in the NSC lattice model (ie. lineages, proportions of cell states and transition rates, cell sizes and number of neighbours, and the aNP-driven LI). To sort out between these hypotheses, we tested whether the NSC lattice model was sufficient to also generate a visible effect of MCs on future MC events: we estimated the Besag’s *L*-function in multiple simulations of the models with and without LI, and quantified the correlation between the position of MCs relative to each other across a range of Δt intervals similar to the in vivo analysis (Figure 7A). Strikingly, we found a prominent statistically significant dispersion of MCs relative to each other with time intervals of Δt4 = 9-12 days in the model including LI compared to a random distribution (Figure 7A). We next showed that this effect statistically differed from the model run without LI, using a permutation test (Hahn, 2012) to compare, with versus without LI, the values of Besag’s *L*-functions quantifying the position of MCs relative to each other at Δt4 time intervals (Figure 7B). Thus, the parameters encoded into the lattice model, and in particular the aNP-driven LI, are sufficient to generate spatiotemporal interactions between MC events that resemble, in their delayed effect and inhibitory output, those detected in vivo.

**Figure 7.**
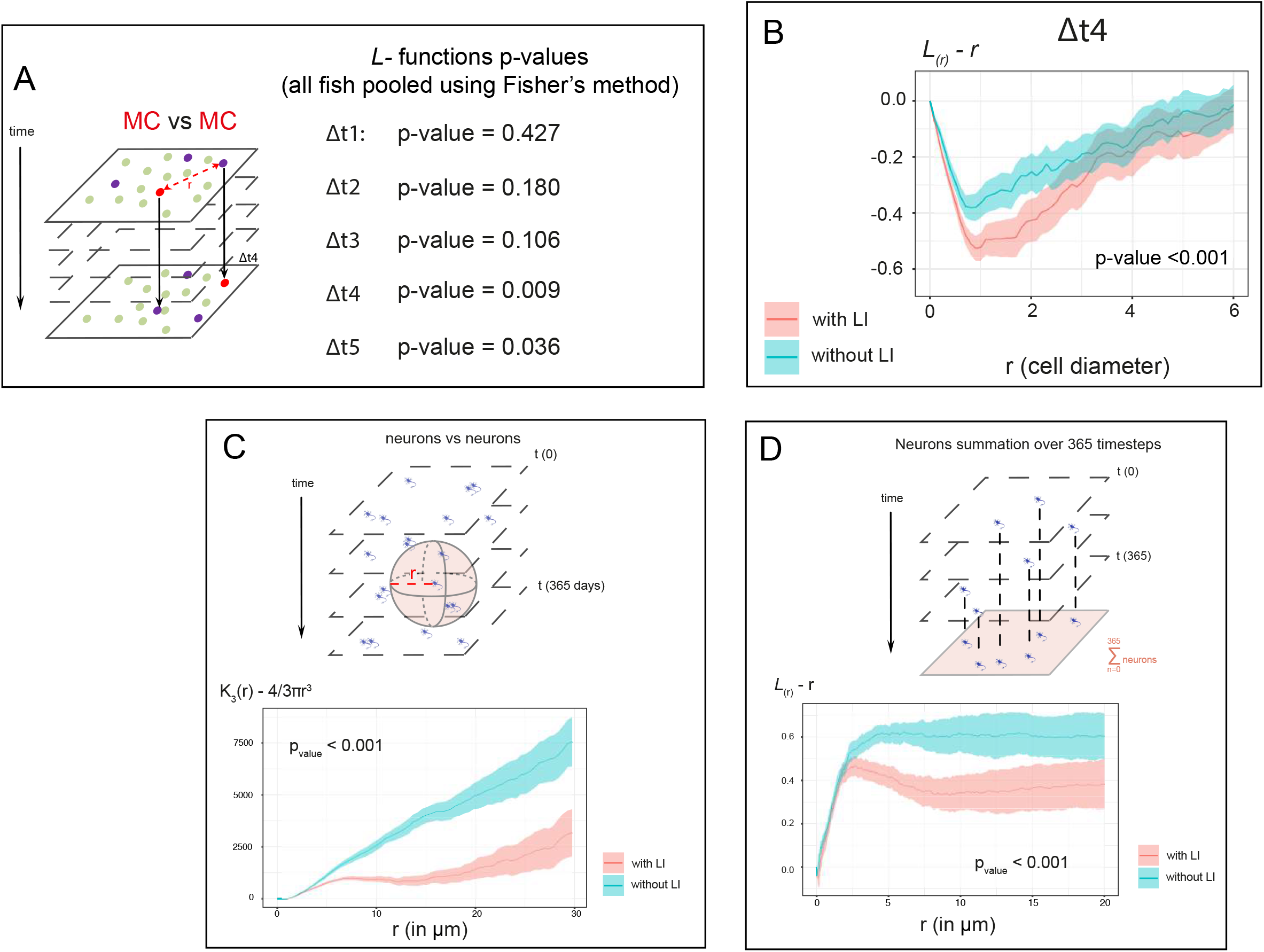
The ‘NSC lattice model’ reproduces the spatiotemporal correlations of NSC fate decisions and shows their impact on long term neuronal distribution. **(A)** Left: MC positions at any time point t in the lattice model are compared using point pattern statistics with MC positions at t+Δt. Right: Combined *p*-values (Fisher’s method) of Besag’s *L*-functions assessing the spatial interaction of MCs relative to each other at all Δt intervals (as in Figure 5) in 18 simulations. A significant *p*-value (0.009) is obtained with Δt4 by combining the results from the 18 simulations. **(B)** Compared *L*-functions (lines) and their 95% confidence intervals (shaded) testing for dispersion between MCs at Δt4 with (pink) and without (blue) aNP-driven LI (18 simulations each). Centered *L*-functions, *L(r) – r*, rather than raw *L*-functions are shown to highlight differences. **(C)** Compared spatial interaction between output neurons with (pink) and without (blue) aNP-driven LI, using a 3-dimensional Ripley’s *K*-function on 18 simulations in each condition, analyzed during 365 time-steps (shaded: confidence intervals). **(D)** Similar comparison in 2D, using the *L*-function. At each time step, the neurons are projected on a 2D plane parallel to the lattice surface. Both **C** and **D** show centered summary functions, *K_3_(r)-(4/3)πr^3^* and *L(r) – r* respectively, rather than raw functions, to highlight differences. *p*-values <0.001: studentized permutation tests comparing the values of the *K*- and *L*-functions.

### The spatiotemporal control of NSC recruitment dynamics supports more homogeneous neurogenesis output

Finally, we aimed to probe the long-term physiological relevance of the uncovered spatiotemporal coordination of NSC dynamics. Two essential characteristics of neurogenesis in the adult pallium are the regular production of neurons along the medio-lateral and antero-posterior axes, and their progressive stacking in age-related layers along the ventro-dorsal axis. These properties generate a homogeneous distribution of neurons in 4D when analyzed over long time frames (Furlan et al., 2017), with, in average, 30μm thick parenchymal layers of neurons generated every 2 months in adults (Than-Trong et al., 2020). To assess the relevance of NSC coordination mechanisms in the long-term generation of this pattern, we ran simulations over 365 days (1 year), and extracted from the model the predicted positions of neurons at all time points. Using a studentized permutation test (Hahn, 2012), the comparison of 3-dimensional Ripley’s *K*-functions between simulations run in the presence versus absence of LI showed that the positions of neurons were significantly less clustered in the former case (Figure 7C). The difference between the two conditions, most notable between ranges 5 μm to 20 μm, is statistically highly significant (*p*-value < 0.001), highlighting that coordinating NSC activation strongly impacts the spatiotemporal distribution of pallial neurons in 3D. The pallium, including Dm, is also composed of distinct neuroanatomical units along the medio-lateral and antero-posterior axes (Ganz et al., 2015). Thus, we also wondered whether the spatiotemporal coordination of NSC activation would additionally impact long-term neuronal distribution in relation with these units. To address this, we reduced the dataset to a 2D dimension by projecting neuronal layers into a monolayer parallel to the dorsal pallial surface. A permutation test (Hahn, 2012) applied to this monolayer revealed that neurons were significantly more clustered under conditions with, versus without, aNP-driven LI (Figure 7D). Together, these results indicate that an important output of the spatiotemporal control of NSC recruitment by population-derived cues is to spatially homogenize neuronal production at long-term across all pallial dimensions, with likely consequences on pallial neuronal identities.

## Discussion

This work addresses the key issue of long-term NSC population homeostasis from the perspective of space and time. Using an interdisciplinary approach, we show that the spatial pattern of NSC activation events responds to, and is temporally propagated by, a combination of local interactions inherent to the NSC pool. Our data provide the first demonstration for the existence of a spatiotemporal coordination of NSC decisions, and identify the qualitative, quantitative, molecular and functional attributes of the lateral inhibition feedback process involved. They also reveal how the transient nature of signaling cells, their downstream position in the NSC lineage and their effect on adjacent cells generate a dynamic and spatiotemporally propagating process that ensures homeostasis of the NSC population at long-term and large scale.

### An “in vivo to modeling” cross-talk to study the spatiotemporal control of NSC decisions in the adult vertebrate brain

Recent studies described a random spatial distribution of NSCs in S phase (Lupperger et al., 2018, 2020), but alternative hypotheses such as local spatiotemporal regulation could not be excluded, and these studies were short-term (72 hours) and merged distinct neuroanatomical domains with reported differences in NSC activation frequencies (Dray et al., 2015). Analyzing the spatiotemporal dynamics of adult NSC populations is challenging: (i) it requires dynamic analyses of cell fate preserving NSC arrangements in their physiological niche, (ii) it involves recording jointly the behavior of each individual NSC and its neighbors, and (iii) it faces the extremely slow dynamics of NSC fate decisions, with quiescence times over months. Intravital imaging methods were recently developed in zebrafish and mouse to film adult NSCs in their niche, and applied to record temporal fate choices in some individual cells (Barbosa et al., 2015; Dray et al., 2015; Pilz et al., 2018; Than-Trong et al., 2020). Here, we made use of the dorsal positioning of the zebrafish adult pallial ventricular zone to access and track the entire NSC population, and to spatially resolve, relative to each other, the state of all NSCs at any time point and of all NSC lineage trees over time. This whole-population tracking (1203 tracks followed) was coupled with spatial statistics to reveal the spatiotemporal parameters and interactions relevant for NSC dynamics, providing the first such analysis in the adult vertebrate brain. The Dm germinal zone encompasses domains homologous to the adult mouse neurogenic niches (reviewed in Labusch et al., 2020). Both species also share lifespans, astroglial NSCs organized in tight pools (März et al., 2010; Mirzadeh et al., 2008; Moss et al., 2016; Seri et al., 2004), equivalent durations of NSC quiescence phases, and compatible NSC clonal dynamics (Basak et al., 2018; Than-Trong et al., 2020; Urbán et al., 2016), stressing the general scope of our findings.

We further exploited this experimental tool to develop a modelling platform based on stochastic simulations on a cell lattice that allows dissecting the spatiotemporal dynamics of NSC activation and division events at steady state. Applying statistical correlation analysis to simulations demonstrates that our model captures both the cellular and the regulatory processes underlying the population dynamics of the adult NSC germinal pool. We show also how this model can be used to elucidate experimental observations, and enables quantitative testing of the origin and output of coordinated NSC behaviors at long term. Together, this “in vivo to modelling” combination now offers unprecedented resolution to unravel NSC population rules and their impact.

### The initial NSC activation event from quiescence is determinant checkpoint that impacts NSC maintenance and output

Our study focuses on two related NSC decisions: activation (ie. cell cycle re-entry from quiescence) and division. We used PCNA expression as the best account of NSC activation in fixed preparations. In contrast, we focused on cytokinesis to refine the temporal window of dynamic analyses, which therefore also read the “activation to division” duration. The distribution rules of NSC division events at each time point in live imaging (Figure 4) are however identical to those describing NSC activation in fixed samples (Figures 1 and 2). Further, by tracking a total of 1203 lineages, the large majority (80%) of activation events were followed by cell division (Figure S5), indicating that the two events are largely slave to one another. Thus, our conclusions more generally apply to NSC recruitment via division. This decision account for 80% of fate choices in the adult zebrafish pallium under physiological conditions (Barbosa et al., 2015; Than-Trong et al., 2020, and this study), and is also a relevant determinant of long-term NSC maintenance (Urbán et al., 2019).

We also exclusively considered de novo activation/division events from quiescence, disregarding the second or third division events of NSCs that rapidly divide repeatedly. This choice is supported by two sets of observations. First, rapid consecutive NSC divisions without intervening quiescence phases would be predicted to generate transiently clustered aNSCs, but these events are rare (14 tracks in a total of 117 dividing tracks followed, ie. 12% of NSCs) (Figure S5). Second, the initial NSC activation decision conditions both neurogenesis and the maintenance of NSC numbers: in both zebrafish and mouse, NSC fates at division are unbalanced, respectively with a high bias towards NSC amplification (Rothenaigner et al., 2011; Than-Trong et al., 2020) or neurogenic fate and loss (Calzolari et al., 2015; Encinas et al., 2011; Fuentealba et al., 2012; Pilz et al., 2018). Thus, NSC activation indirectly impacts the quantitative distribution of fates. Our findings therefore also provide a first mechanistic insight into the intrinsic coordination processes that quantitatively balance the maintenance and output of adult NSC niches over time.

### Lineage-derived inhibitory feedbacks generate an intrinsic niche for NSC coordination

A major finding of our study is the demonstration that NSC activation events are spatiotemporally controlled by cell-cell interactions operating between progenitors, putting forward the concept of an “intrinsic niche”. This contrasts with situations where SC properties depend on local external niches, as in the adult intestine, stomach, colon or hair follicle: there, SC identity and/or fate is also flexible but spatially biased relative to local external cues such as geometry or non-progenitor cell types (Gehart and Clevers, 2019; Ritsma et al., 2014; Rompolas et al., 2012). In an intrinsic niche, SC population homeostasis is both the consequence and the origin of an internal systems dynamics. While the two principles are not mutually exclusive, an intrinsic niche is perhaps particularly suitable in the context of large SC populations that need to homogeneously generate progeny cells over extended distances, such as NSC niches or the interfollicular SC population in the adult mammalian skin (Belokhvostova et al., 2018; Blanpain and Fuchs, 2009). Interestingly, in the latter case, a recent study using intravital imaging showed that neighboring SCs coordinate their fate decisions to differentiate and divide (Mesa et al., 2018).

Our work also identifies major cellular actors of the intrinsic niche of the adult pallium: aNPs and dividing NSCs (MCs). aNPs appear as key systems components that decrease the propensity for NSC activation in their immediate vicinity at any time, by 2-to 4-fold (Figures 2 and S3). MCs, directly or indirectly, have a similar effect with a temporal delay of 9-12 days -although the strength of this inhibition could not be evaluated- (Figure 5). It remains to be precisely identified what is dynamically read as the effect of MCs: this may be cytokinesis per se or associated changes of cell morphology, activation, or any other obligatory associated event such as lineage progression and the generation of aNPs themselves. Our modeling results point to aNPs as important components of the regulation exerted by MCs. Indeed, encoding the inhibitory effect of aNPs in the model appears sufficient for MCs to impact future neighboring MCs with a temporal delay similar to that measured in vivo. Thus, the effect of MCs is likely linked with lineage progression and aNP generation, although additional mechanisms cannot be formally excluded.

The lineage relationship existing between receiving (qNSCs) and signaling cells (MCs and aNPs), further indicates that one coordination rule of the intrinsic niche is akin to a retroactive control. Such a process, gated via its own output, was theoretically modeled to instruct balanced cell fate choices within homeostatic cell populations (Lander et al., 2009), although without a spatial component. The mechanism unraveled here is akin to a lateral inhibition process, well known in many developmental systems, where it is frequently driven by Notch signaling (Moore and Alexandre, 2020; Sjöqvist and Andersson, 2019). In the adult zebrafish and mouse, NSCs are maintained in quiescence by Notch2/3 signaling (Alunni et al., 2013; Chapouton et al., 2010; Ehm et al., 2010; Engler et al., 2018; Imayoshi et al., 2010; Kawai et al., 2017). In the SEZ, this involves DLL1-mediated signaling from aNSCs and/or aNP equivalents (Ables et al., 2010; Kawaguchi et al., 2013). Our work importantly extends these findings by highlighting the spatiotemporal relevance of this mechanism in adult NSC niches. It also quantifies its properties and dynamics, and provides evidence for its spatiotemporal impact on neurogenesis output. It is to note that only 64% of NSC divisions (asymmetric and neurogenic) lead to the production of aNPs (Than-Trong et al., 2020), and that the majority of NSCs at any time point are not in contact with aNPs (Figure S6F). Our in vivo and modeling results however suggest that the effect of aNPs can be sufficiently strong to be visible at a global scale on NSC activation events. It remains possible that, in vivo, aNSCs, many of which express DeltaA, act as an additional source of Notch-mediated feedback. It will be important to directly measure the spatiotemporal dynamics of Notch3 signaling activity in situ in relation with candidate signaling cells.

### The adult pallial intrinsic niche permits the dynamic, self-sustained and self-propagating coordination of NSC activation events in time and space

Another important advance of our work is the identification of several of the fundamental systems properties of the adult pallium intrinsic niche, with respect to NSC activation control.

First, the interactions coordinating NSC state are transient. This is due both to the short half-life of signaling cells and to the labile state that they encode -quiescence-. Our NSC lattice model is key to interpret the bases and output of such interactions, showing that they are sufficient to generate balanced dynamics and maintain system’s equilibrium and spatiotemporal correlations over durations equivalent to a lifetime. The fact that the process never becomes fixed stands in contrast with a number of lateral inhibition-mediated processes during embryogenesis, for example alternative fate choices or self-organized pattern generation such as in the Drosophila notum or vertebrate inner ear, where final cellular states are encoded (Brown and Groves, 2020; Schweisguth and Corson, 2019). We propose that this is particularly relevant in the case of adult SC ensembles, which need to maintain homeostasis over a lifetime while accommodating lineage generation. Second, these interactions operate with a temporal delay. As the NSC lattice model suggests, this delay is likely due, at least in part, to lineage progression from the aNSC to the aNP state, but may additionally involve other mechanisms, yet to be discovered. Typically, when a quiescence-promoting signal such as Notch3 signaling is abolished, or when NSC activation is triggered for repair, NSCs do not activate immediately (Alunni et al., 2013; Baumgart et al., 2012; März et al., 2011).

Physiologically, the duration or depth of G0 or the different checkpoints needed for activation may thus introduce a delayed output from the moment when an inhibitory signal, eg. exerted by a neighboring MC, is released.

Third, because the inhibitory interactions that we uncovered are exerted on neighbors, these interactions control the spatial distribution of cell states. This is a fundamental point, as in the zebrafish pallium the absence of extensive cell migration implies that NSC activity determines neuronal location (Furlan et al., 2017). Indeed, our NSC lattice model shows that the coordination of NSC states via aNP-driven LI decreases neuronal clustering over a lifetime, hence is important for long-term pallial architecture, and presumably function. Likewise, in neurogenic niches in the adult mouse, SGZ neurons remain local, and NSC distribution within the SEZ biases neuronal identity (Fiorelli et al., 2015; Gonçalves et al., 2016; Obernier and Alvarez-Buylla, 2019).

In line with these properties, the NSC lattice model is unique: First, unlike mathematical models of embryonic lateral inhibition (Binshtok and Sprinzak, 2018), it captures the spatiotemporal properties of a dynamic homeostatic system rather than a fixed final pattern. Second, it describes a system where cell morphology is coupled with cell state, a feature which is important for proper spatial organization. Thus, it provides a new set of tools allowing theoretical analyses, particularly suited for spatiotemporally coordinated and dynamic SC populations.

Together, the systems parameters that we uncovered transform local and short-term interactions into large-scale and long-term coordination that homogenize NSC behavior across the niche during a lifetime. Interactions between SCs are starting to be identified as important components of SC fate in several systems, revealing other intrinsic niches. In the interfollicular epidermis of the adult mouse, SC differentiation and division events are spatiotemporally coupled, with a delay of 12 hours and following the transmission of a mechanical signal to neighboring SCs (Mesa et al., 2018). Likewise, SC loss triggers neighboring SC division in the Drosophila gut (Liang et al., 2017; De Navascués et al., 2012). Although the rapid SC turnover of these systems strongly contrasts with the slow dynamics of adult vertebrate NSC niches, and the signaling mechanisms may be different, all illustrate a common principle where SC states and spatial organization are maintained stable at a local scale through spatiotemporal interactions that then gradually propagate through the whole germinal tissue. We propose that such mechanisms may more generally define the principles of intrinsic niches in adult SC systems.

## Supporting information

Supplementary figures, tables and discussion

## Acknowledgements

We thank members of the ZEN team for their constant input, and Boris Shraiman (UC Santa Barbara) and Micha Hersch (University of Lausanne) for providing Matlab functions that helped generate random cell lattices and execute minimization of the lattice mechanical energy. Funding: Work in the L. B-C. laboratory was funded by the ANR (Labex Revive), La Ligue Nationale Contre le Cancer, Centre National de la Recherche Scientifique, Institut Pasteur, DIM ELICIT (together with E.B.) and the European Research Council (AdG 322936). L.M. was recipient of a PhD student fellowship from Labex Revive and the Fondation pour la Recherche Médicale (FRM). D.S. and U.B. acknowledge the support of the European Research Council (ERC) under the European Union’s Horizon 2020 research and innovation programme (Grant agreement No. 682161). U.B. is recipient of a fellowship from The Marian Gertner Institute for Medical Nanosystems at Tel Aviv University. Multiphoton equipment in E.B.’s laboratory was partly supported by Agence Nationale de la Recherche (ANR-11-EQPX-0029 Morphoscope2, ANR-10-INSB-04 France BioImaging).

## Authors contribution

N.D. generated all intravital imaging data, with technical advice from S.B., P.M. and E.B.; N.D. and L.M. generated and analyzed all fixed biological preparations; S.B. and S.O. prepared all animals for imaging; M.K. and E.B. provided the proof-of-principle evidence of the do-ability of intravital imaging; W.S. conducted the point pattern analyses that revealed cell interactions at an initial stage of the work; U.B. developed the mathematical model of NSC behavior under the supervision of D.S.; S.H., J-B.M. and J-Y.T. worked on statistical methods quantifying interactions in individual samples; F.C. conducted all final spatial statistics analyses of experimental data and simulations, under the supervision of G.L.; N.D. and L.B-C. designed the study and wrote the manuscript, with input from all authors.

## Declaration of interests

The authors declare no competing interests.

## STAR * METHODS

### KEY RESOURCES TABLE

| REAGENT or RESOURCE | SOURCE | IDENTIFIER |
| --- | --- | --- |
| Antibodies |  |  |
| Mouse monoclonal (IgG1) anti- ZO1 | Thermo Fisher | Cat#33-9100, RRID: AB_2533147 |
| Mouse monoclonal (IgG1) anti- SOX2 | Abcam | Cat#ab171380, RRID: AB_2732072 |
| Rabbit anti-DsRed | Takara | Cat# 632496, RRID:AB_10013483 |
| Rat monoclonal anti-BrdU | Abcam | Cat#ab6326; RRID: AB_305426 |
| Chicken anti-GFP Antibody | Aves lab | Cat#GFP-1020; RRID: AB_10000240 |
| Rabbit Polyclonal anti-PCNA | GeneTex | Cat#GTX124496; RRID: AB_11161916 |
| Mouse anti-PCNA IgG2a (PC10) | Santa Cruz Biotechnology | Cat# sc-56, RRID:AB_628110) |
| Goat anti-Mouse IgG2a Alexa633 conjugated | Thermo Fisher Scientific | Cat# A-21136, RRID:AB_2535775 |
| Goat anti-Mouse IgG1 Alexa546 conjugated | Thermo Fisher Scientific | Cat# A-21123, RRID:AB_141592) |
| Goat Anti-Chicken IgG(H+L) Alexa488 Conjugated | Thermo Fisher Scientific | Cat# A-11039, RRID:AB_142924 |
| Goat Anti-Rabbit IgG(H+L) Alexa405 Conjugated | Thermo Fisher Scientific | Cat# A-31556, RRID:AB_221605 |
| Goat anti-Rat IgG-488 IgG(H+L) Alexa488 Conjugated | Thermo Fisher Scientific | Cat#A-11006, RRID:AB_2534074 |
| Chemicals, Peptides, and Recombinant Proteins |  |  |
| LY-411575 | Sigma-Aldrich | Cat#SML0506 |
| 5-Bromo-2-deoxyuridine (BrdU) | Sigma-Aldrich | Cat#B5002 |
| Antigen Retrieval (HistoVT One) | Nacalai Tesque | Cat#06380-05 |
| Deposited Data |  |  |
| Experimental Models: Organisms/Strains |  |  |
| Zebrafish: AB WT strain | N/A | ZFIN: ZDB-GENO-960809-7 |
| Zebrafish: Tg(*gfap:GFP*)mi2001 | Zebrafish International ResourceCenter, Eugene, OR | ZDB-TGCONSTRCT-070117-154 |
| Zebrafish Tg(*mcm5:EGFP*) | Zebrafish International ResourceCenter, Eugene, OR | ZFIN ID: ZDB-TGCONSTRCT-160126-1 |
| Casper (*roy-/-;nacre-/-*) (*mitfaw2/w2; mpv17a9/a9*) | Zebrafish International ResourceCenter, Eugene, OR | ZDB-FISH-150901-6638 |
| Zebrafish Tg(*deltaA:GFP*) | Zebrafish International ResourceCenter, Eugene, OR | ZFIN ID: ZDB-TGCONSTRCT-100823-1 |
| Software and Algorithms |  |  |
| Black Zen software | Carl Zeiss | RRID:SCR_018163 |
| Imaris | Bitplane | http://www.bitplane.com/Imaris/Imaris,<br>RRID: SCR_007370 |
| Fiji | Fiji | http://fiji.sc, RRID:<br>SCR_002285 |
| Matlab 2017a | Matlab | MATLAB,<br>RRID:SCR_001622 |
| R Project for Statistical Computing | R Project for Statistical Computing | http://www.r-project.org/,<br>RRID:SCR_001905 |
| Prism 8 | GraphPad | http://www.graphpad.com/,<br>RRID:SCR_002798 |
| Illustrator CS6 | Adobe | Adobe Illustrator,<br>RRID:SCR_010279 |
| Other |  |  |

### LEAD CONTACT AND MATERIALS AVAILABILITY

Further information and requests for resources and reagents should be directed to and will be fulfilled by the Lead Contact, Laure Bally-Cuif.

### EXPERIMENTAL MODEL AND SUBJECT DETAILS

#### Fish husbandry and lines

All animal experiments were carried out in accordance to the official regulatory standards of the department of Essonne (agreement number A 91-577 to L.B.-C.) and department of Paris (agreement number A 75-1522 to L.B.-C. and N.D.) and conformed to French and European ethical and animal welfare directives (project authorization from the Ministère de l’Enseignement Supérieur, de la Recherche et de l’Innovation to L.B.-C.). Zebrafish were kept in 3.5-liter tanks at a maximal density of five per litter, in 28.5°C and pH 7.4 water. Three-to four-month-old adult zebrafish were used; *Tg(gfap:eGFP)* (Bernardos and Raymond, 2006), *Tg(gfap:dTomato)* (Satou et al., 2012), *Tg(mcm5:eGFP)*^gy2^ (Dray et al., 2015) and Tg(*deltaA:GFP*) (Madelaine and Blader, 2011). For live imaging Tg(*gfap:dTomato*)/+;Tg(*mcm5:eGFP*)^gy2^/+ in the *casper* double mutant background (*roy^−/−^;nacre^−/−^*) (White et al., 2008) were used.

### METHOD DETAILS

#### BrdU pulse labeling

BrdU was applied in the fish swimming water for 3h at a final concentration of 1 mM for BrdU. After the pulse, fish were transferred to a tank with fresh fish water during chase period of 24h.

#### Filtering PCNA signals for pre-cytokinesis events

To define the criteria for this filtering, we applied a BrdU pulse to 3mpf animals, followed by a short chase time sufficient for a cell cycle (24 hours) (Alunni et al., 2013), and measured PCNA in BrdU^+^ doublets (Figures S2A-C). BrdU^+^;GFAP^+^ doublets were rarely observed at chase time 0 (8% of cases) (not shown), thus neighboring aNSCs found synchronously in S phase are very rare events. At 24 hours post-BrdU, 97% of BrdU^+^;GFAP^+^ cells were found as doublets, with both cells PCNA^+^ in 80% of these doublets (Figure S2C). Thus, at any given time, the large majority of GFAP^+^;PCNA^+^ cells found as adjacent pairs are post-cytokinesis sister NSCs, captured within the 24 hours following division. A morphological analysis at high resolution further indicated that such sister cells remain tightly juxtaposed and display generally symmetrical apical domains (Figure S2D). On this basis, we manually filtered the map to replace doublets of adjacent and morphologically symmetrical GFAP^+^;PCNA^+^ cells -representing post-cytokinesis events-by an equivalent doublet of qNSCs (Figure S2E-G, Table S1).

#### LY411575 treatment

Stock solutions of LY411575 (LY) at 10 mM were prepared by disolving 5 mg of LY in 1.05 mL DMSO and stored at −80°C until use. To block Notch signaling, LY was applied in the fish swimming water at a final concentration of 10 mM (Alunni et al., 2013). The solution was refreshed every 24 h. Control fish were treated with the same final concentration (0.1%) of DMSO carrier.

#### Whole-mount Immunohistochemistry

Brains were dissected in PBS, transferred to a 4% paraformaldehyde solution in PBS for fixation (2 hours at room temperature or overnight at 4°C) dehydrated and kept in 100% methanol at −20°C. Brains stored in 100% MeOH were rehydrated and washed 3 times with PBST (0.1% Tween-20 in PBS). For BrdU an antigen retrieval step was performed with an incubation in 2 M HCl (Sigma-Aldrich, 258148) at room temperature for 30 min. For proliferating cell nuclear antigen (PCNA) immunolabeling, an antigen retrieval step was performed with an incubation in HistoVT One (Nacalai Tesque) for an hour at 65°C. Brains were then washed three times for 5 min each with PBST, incubated into Blocking Solution (5% Normal Goat Serum, 0.1% DMSO, 0.1% Triton X-100 in PBS) (Sigma Life Science, 1002135493) for 1 h at RT. Primary antibodies were diluted in blocking Solution and incubated for 24 h at 4°C. The following primary antibodies were used: anti-ZO1(Mouse, 1:200, Thermo Fisher Scientific), anti-SOX2 (Mouse, 1:200, Abcam), anti-DsRed (Rabbit, 1:250, Takara), anti-GFP (Chicken, 1:500, Aves Labs), anti-BrdU (Rat, 1:150, Abcam), anti-PCNA (Rabbit, 1:500, GeneTex), anti-PCNA (Mouse, 1:500, Santa Cruz Biotechnology). Brains were subsequently washed six times for 15 min with PBST and incubated for 24h at 4°C with secondary antibodies diluted 1:1000 in Blocking Solution.

The following secondary antibodies were used: anti-IgG2a conjugated to Alexa633 (Thermo Fisher Scientific), anti-IgG1 conjugated to Alexa546 (Thermo Fisher Scientific), anti-Chicken IgG(H+L) conjugated to Alexa488 (Thermo Fisher Scientific), anti-Rabbit IgG(H+L) conjugated to Alexa405 (Thermo Fisher Scientific), anti-Rat IgG conjugated to Alexa488 (Thermo Fisher Scientific). Brains were then washed six times for 15 min with PBST.

#### Confocal imaging whole-mounted immunohistochemistry

Fluorescent images of whole-mount telencephali were acquired on confocal microscopes (LSM700 and LSM710, Zeiss), using a 40X oil objective (Plan-Apochromat 40x/1.3 Oil M27) with an optical sectioning in Z every 0.5μm and a tile scan of 4 to 8 Z-stacks. Stitching was done with the ZEN software after imaging.

#### Fish anesthesia and mounting for intravital imaging

Anesthesia and mounting for imaging were conducted as in previous studies (Dray et al., 2015; Than-Trong et al., 2020). Briefly, anesthesia was initiated by soaking the fish for approximatively 90 s in water containing 0.01% MS222 (Sigma-Aldrich). They were then transferred into a water solution of 0.005% (v/v) MS222 and 0.005% (v/v) isoflurane to maintain the anesthesia during the whole duration of the imaging session and mounted in a home-made plastic dish between pieces of sponge. Overall, fish were anesthetized for about 30 min per session.

#### Multiphoton intravital imaging and analysis

The intravital imaging was performed on a customized commercial multiphoton microscope (TriM Scope II, LaVision BioTec) equipped with an ultrafast oscillator (λ = 690 to 1300 nm; InSight DS+ from Spectra-Physics Newport) and a MaiTai laser (TI:Sapphire, λ = 690–1040 nm Spectra-Physics). dTomato was excited at 1120 nm and GFP at 950nm. The fluorescent signal was collected with GaAsP detector (H7422-40, Hamamatsu). To image the entire volume of interest, spanning typically 800 μm by 800 μm by 250 μm (i.e., a single brain hemisphere), we recorded mosaics consisting of four z-stacks with an overlap of 10%. For each z-stack, the lateral field of view was 405 μm by 405 μm, the depth of imaging varied from 250 to 290 μm (starting about 250 μm below the skin), the voxel size was 0.8 μm by 0.8 μm by 2 μm, and the pixel dwell time was 4.9 μs.

### QUANTIFICATION AND STATISTICAL ANALYSIS

#### Image analysis of immunohistochemistry

3D renderings were generated using the Imaris^®^ software (versions 8 and 9, Bitplane). The 3D image was cropped to feature only the pallium as our region of interest and histograms was adjusted for each channel. The images were segmented manually using semi-automatic detection with the Imaris spots function followed by manual curation in the dorsomedial part (Dm) of the pallium (Dray et al., 2015; Than-Trong et al., 2020). Except for the BrdU experiment, at least 3 brains were analyzed per condition (WT, LY, DMSO), n represent the number of brains analyzed (one hemisphere per brain with 700 to 1919 cells counted per hemisphere), SEM and SD are presented. For the BrdU experiment 2 fish were analyzed and n represents the total number of cells. Mean cell diameters (r=1) are the mean distances between qNSCs, aNSCs and aNPs (Figure 1E and F: 9.92μm; Figure 2D: LY: 11.21μm, DMSO: 9.71μm; Figure S3: mean 10.07μm, SD 0.35; Figure S4: mean LY: 11.02μm, SD 0.30, mean DMSO: 11.06μm, SD 1.2), qNSCs and aNSCs or qNSCs, aNSCs and MCs (Figures 4 and 5, respectively) (Bibi: 12.88μm, Mimi: 13.19μm, Titi: 13.79μm), measured with the Imaris^®^ software. In all spatial analyses of simulations, mean cell diameters (r=1) are mean distances between qNSCs, aNSCs and aNPs (Figures 6E”, 6E’’’ and 7B: average cell diameter over all the simulations and timesteps, 10μm; Figures 7C and 7D: we converted the simulated units into microns by comparing the average cell diameter from the simulation to the one from the experiments [10μm for the x-y plane, see Figure S3] and by using the experimental stacking rate of neurons (this is 30μm over 2 months for the z axis) (Than-Trong et al., 2020).

Statistical analyses (except the spatial statistics) were carried out using Prism and Microsoft Excel. All the statistical tests performed were two-tailed, and their significance level was set at 5%.

#### Image analysis of intravital imaging

Image were combined and analyzed as in previous studies (Dray et al. 2015). Briefly, Z-stacks acquired on successive imaging were first converted into a single file after cropping two files in the three dimensions using Imaris^®^ (Bitplane) or Fiji. The alignment was done at the cellular level using landmark-based registration for which a few cells are detected and manually tracked over time and their average drift was corrected using Imaris. The histograms of fluorescence intensity was adjusted ‘by eye’ (linear stretch of the histograms) to correct the minor fluctuations in intensity from one day to another. After alignment, all cells were manually detected using Imaris^®^ and their position where exported for further analysis via Matlab^®^. Three fish were analyzed (mimi, titi and bibi), n represent the number of brains analyzed (one hemisphere per brain) with 300 to 500 cells tracked over eight time points every 3 to 4 days over 23 days total (one to two more time points were acquired after these 23 days by not analyzed because the time interval was longer. Statistical analyses (except the spatial statistics) were carried out using Prism and Microsoft Excel.

#### Spatial statistics (Point pattern analysis)

##### Summary statistics

To determine whether there exists an interaction between two specific states of cells *i* and *j*(*i, j* = aNSC or aNP), we analyse their relative positions using three multitype second-order summary statistics (Figure S3):

- Besag’s *L*-function (Besag, 1977) is a variance-stabilised version of Ripley’s *K*-function (Ripley, 1977) which averages the number of cells of state *j* within a distance *r* of a typical cell of state *i*. It is a popular technique for analysing spatial correlation in point patterns, usually by visually inspecting the empirical 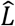-function, calculated from the data, and comparing it to the theoretical *L*-function of the homogeneous Poisson process *L_pois_*(*r*) = *r*, which serves as a benchmark for ‘no correlation’: for example, if 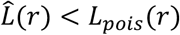, this indicates that there are fewer neighbors than would be expected for a completely random pattern, consistent with some type of inhibition or repulsion between cells. While the *L*-function is good at detecting spatial correlation, it is not adapted to measure the range of said interaction, as it is of cumulative nature: for example, with patterns exhibiting inhibition at small spatial scales, the cumulative effect could still be visible at larger spatial scales.
- The pair correlation function *g* counts contributions from cells of state *j* at a distance equal to *r* of a typical cell of state *i*: it is the derivative of the *K*-function, rescaled such that, for a typical random process, *g_pois_*(*r*) = 1. As a complementary tool to the *L*-function, the *g*-function helps detect the range of interaction between cells: an empirical value *ĝ*(*r*) ≠ 1 suggests that there is a spatial correlation between cells distant of *r*. Both the *L*- and *g*-functions must be corrected for edge effects to avoid the bias that occurs when counting the number of neighbors for cells close to the border of the observed area. In this paper, we use an isotropic correction to account for edge effects. Details on the theoretical properties, statistical estimation, computation and edge corrections of the *L*- and *g*-functions be found in (Baddeley et al., 2016).
- The *M*-function has been introduced recently in (Marcon and Puech, 2010; Marcon et al., 2012), as an extension of the *K*-function. It is of cumulative nature and measures the frequency of cells ot state *j* within a distance *r* of cells of state *i*, relative to that over the whole observed area. One of its advantages is that it is easily interpreted: for example, 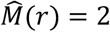 would indicate that the relative density is double that in the observed area.

##### Hypothesis testing and simulation envelopes

To assess whether the interaction between cells of states *i* and *j* is statistically significant, we perform Monte Carlo tests based on simulation envelopes of the summary functions. These envelopes are constructed by generating simulated patterns and their summary statistics from the “random labelling” null hypothesis 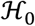, which assumes that the state of any cell is random with fixed probabilities and independent of other cells’ positions. The simulated patterns are generated by randomly permuting cells of state *j* amongst all cells while keeping those of state *i* fixed. The simulation envelopes provide acceptance intervals (or non-rejection intervals) for the null hypothesis: if the empirical function lies outside the simulation envelopes, then 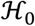 may be rejected at a significance level that is inversely proportional to the number of patterns simulated (Figure S3A). Note that 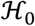 is different from the usual “complete spatial randomness and independence” null hypothesis, that assumes that the underlying processes for cells of state *i* and *j* are independent Poisson patterns, which is not a good model for cell patterns, as cells are subject to the hardcore constraint (no two cells can be too close to each other). In particular, the aforementioned empirical summary functions should not be compared to the theoretical value for the Poisson pattern.

We consider two types of envelopes, depending on whether the interaction is tested at a specific distance *r* or over a range [0, *R*] of distances:

- Simultaneous envelopes are constructed by measuring, for each simulated pattern, the most extreme deviation between the empirical and expected values of the summary statistic, where the maximum deviation is taken over the range [0, *R*], They correspond to a global test of spatial correlation, indicating that an interaction exists for cells within a distance *R* if the observed curve ever lies, at any distance *r* < *R*, outside the simulation envelopes. We consider simultaneous envelopes for the *L*-function only, as it is the only function considered that has a stable variance with respect to *r*. A second strategy for detecting spatial correlation for the *L*-function is the Diggle-Cressie-Loosmore-Ford (DCLF) test, as proposed by (Cressie, 1991; Diggle, 1986; Loosmore and Ford, 2006). Instead of the most extreme deviation, the test uses the integrated squared deviation between the empirical and expected values of the summary statistic over the range [0, *R*]. It is complementary to the maximum deviation envelopes to detect spatial interaction between cells.
- Pointwise envelopes are constructed by measuring, for each simulated pattern and each distance r, the most extreme deviation between the empirical and expected values of the summary statistic. Their interpretation requires the distance *r* to be set in advance of the analysis. We consider pointwise envelopes for the *g*-function only, as a way to detect the range of spatial interaction if it exists (*i.e*. if the global test on the *L*-function has rejected 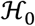). Ranges considered are *r* = 1,2 and 3 average cell diameters. Details on hypothesis tests and simulation envelopes for point patterns can be found in (Baddeley et al., 2016).

##### Statistical approach

In view of the preceding methods, we consider the following statistical approach for the analysis of interaction between cells of states *i* and *j* on a homogeneous part of the brain (the Dm domain): *(i)* first, we consider global tests (both maximum deviation and DCLF) for the *L*-function: this indicates whether an interaction exists (Figures 1E and 2D); *(ii)* then, if an interaction exists, we consider pointwise tests for the *g*-function at distances *r* = 1,2 and 3 average cell diameters: this indicates the exact range of the interaction (Figure 1F); *(iii)* finally, we measure the strength of the interaction with the *M*-function (Figure 1F).

In order to determine whether the interaction between aNP and aNSC cells exists beyond the first-order neighbors (those cells that share a membrane), we calculate the distance between a typical aNP cell and its furthest first-order neighbor (which, for aNP cells, is in average the fourth nearest neighbor, since they have four direct neighboring cells). aNP cells too close to edges were excluded from this calculation, to avoid evident bias. We then report the ninety-fifth 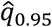 centiles for this distance on the graph of the *g*-function (vertical green line) (Figure 1F). If the empirical 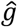-function does not lie outside the pointwise envelopes beyond this range, then it is unlikely that the interaction goes beyond direct neighboring cells.

##### Dynamic patterns

To determine whether some inhibition also occurs between division events, we consider the patterns at time steps t and t+Δt and note state i the mother cells of the former pattern and j those of the latter. In order to include the whole observation period, the L-, g- and M-functions for each pair of patterns t and t+Δt (t=1,…,T-Δt) are pooled using a weighted average (Baddeley et al., 2016). Then, the same statistical approach as above can be employed to determine whether an interaction exists between mother cells (Figures 5 and 7).

Note that one-sided rather than two-sided statistical tests are used, assessing only whether dispersion between division events exists, but not clustering, as this analysis was motivated by the previous which found a statistically significant dispersion exerted by aNPs on aNSCs. Consequently, simulation envelopes are one-sided, and the DCLF test only takes into account the negative part of the deviation, i.e. only focuses on the ranges where the empirical summary function is below the expected one.

Some inhomogeneity in the density of cells of dynamic patterns was present due to the curvature of the germinal layer, manifesting as a gradient along the length of the live samples. To account for this inhomogeneity, the position of cells along this dimension was first rescaled according to a smoothing estimate of the density, before running the aforementioned spatial analysis.

Finally, to combine the results from the three fish, we use Fisher’s method (Fisher, 1954; Mosteller and Fisher, 1948) to pool the *p*-values *p_i_* from each hypothesis test into one test statistic:

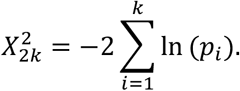

Under the null hypothesis, these *p*-values follow a uniform distribution on the interval [0,1], and hence 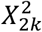 follows a chi-squared distribution with 2*k* degrees of freedom (Figures 5 and 7).

##### Analysis of simulations from the NSC lattice model

To ascertain whether the model correctly mimics the live patterns, we performed a similar spatial analysis of the model simulations, with and without LI. For the comparison of NSCs, summary functions were computed on days 300, 350, 400, 450 and 500, then pooled for each simulation (Figure 6). For the comparison of MCs, since in live patterns time steps were observed three days apart, with MCs in a time step corresponding to all the division events since the previous time step, the analysis of the model simulations was similarly set up: the MCs in every three consecutive days were grouped into one pattern, then each pair of these patterns t and t+Δt was analysed then pooled in the same way as live patterns. Finally, for any given Δt, the results from the simulations were combined using Fisher’s method to pool their p-values (Figure 7).

The comparisons between the situations with versus without LI, of the interactions between MCs (Figure 7B) and between neurons (Figure 7C and D) were carried out using permutation tests proposed by (Hahn, 2012), which consists in comparing the expected summary statistics for each situation. Formally, the test statistic is the integrated squared normalized difference between the expected L-functions with and without LI. The Monte-Carlo p-value of the test is then evaluated by comparing the observed data with random permutations of the L-functions among the groups. Neurons were both analyzed in 3D (Figure 7C), and in 2D (Figure 7D) by projecting their positions to the plane parallel to the pallial surface. For the three-dimensional analysis of neurons, the summary function used is instead Ripley’s K-function, since the three-dimensional analogue to the L-function is not variance-stabilised (Baddeley et al., 2016).

##### See also Supplementary discussion on Statistics

##### Code reproducibility

We used R (R Core Team 2020) software, and packages spatstats (Baddeley and Turner, 2005) and dbmss (Marcon et al., 2015) to do the spatial and spatiotemporal analyses of NSCs, division events and neurons. The code and the datasets used in the publication are publicly available at https://github.com/fcheysson/zebrafish-project.

#### Generation of the NSC lattice model: mathematical approach

To gain insight into the dynamics of NSC activation, division, and differentiation, we developed a statistical spatiotemporal model of NSC behavior (workflow in Figure S7A). The model contains two layers: (i) A steady state mean field analysis of dynamic rate equations describing the transitions between the different progenitor cell states – qNSCs, aNSCs, aNPs, and differentiated neurons. (ii) The ‘NSC lattice model’ which contains stochastic simulations based on a 2D vertex modeling platform that simulate transition between cell states, cell divisions, and cellular differentiation in a disordered cell lattice (Figure 6B).

##### Analytical model

We wanted to develop an analytical model that captures the NSC cell fate transitions (Figure 1C), with the experimentally observed cell fate proportions (Figures 1D and S1D), transition rates (Figure S6B), and considers the inhibition of qNSC activation by neighboring aNPs (Figure 1E-F). The possible transitions considered in the model are shown in Figure 6A.

##### Dynamic equations

The variables in our model correspond to the four different cell states:

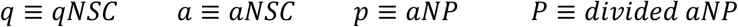

We define ‘divided aNP’ as a different cell state since we have made a simplifying assumption that aNP can divide only once before differentiating into neurons, namely *p* can divide to two *P*. This is based on the experimental observation that clusters of three or more aNPs are rare, compared to singlets or doublets (Figure S6E).

We also denote the transition rates and the differentiation probabilities following cell divisions in the following way:

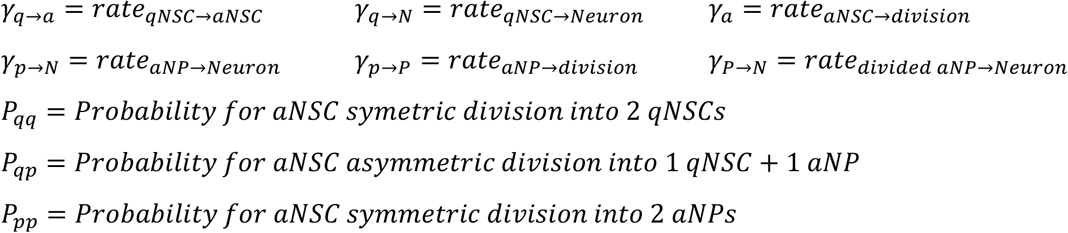

The differential equations corresponding to the 4 cell states are (Figure 6A):

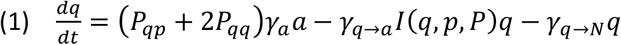

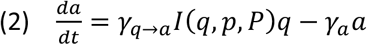

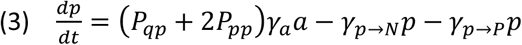

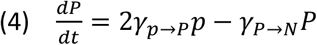

Here, the lateral inhibition (LI) on qNSC transition rate into aNSC, by a neighbor aNP, is written as a multiplication between the parameter *γ_q→a_* and a function *I*(*q,p,P*) which depends on *q, p* and *P*, and satisfies *I* ∈ (0,1]. Depending on neighboring qNSC and aNP, *I* = 1 in case a qNSC does not neighbor an aNP, otherwise *I* < 1. Thus, *γ_q→a_* is the rate for *qNSC* → *aNSC* in case a qNSC does not neighbor an aNP and therefore it is not inhibited. We note that our model does not consider a situation where aNSC transitions back to a qNSC state without dividing since these are rarely observed experimentally.

##### Steady state and mean field analysis

It is understood that different qNSCs will have a different transition rate due to LI that depends on time and space (e.g. on the presence or absence of aNP neighboring cells). In the following steady state analysis we replace the many possible interactions between *qWSCs* and *aNPs* by an approximated mean value for *I*. Thus, at steady state, we get a roughly constant number of cells that are inhibited and therefore a constant average reduction factor denoted *I*\*:

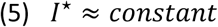

Accordingly, *γ_q→a_I*\* is an average reduced transition rate. For example, if on average half the qNSC cells are completely inhibited from making a q->a transition, then 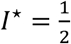 and the average transition rate is 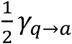. Such approximation is known as a mean field approximation where in the above described system the LI effect is treated as an effective external field rather than a multi-cell-to-cell interactions. We ultimately use this approximation, along with measured experimental values, to extract the dynamic parameters from the model.

At steady state, equations (1), (2), (3) and (4) equal to zero and we get the following relations:

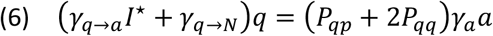

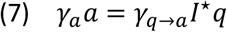

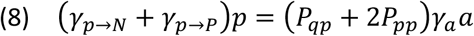

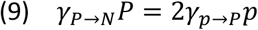

Which leads to a condition for a steady state:

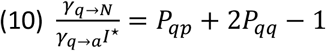

From equation (10) one can see that *qNSC* direct differentiation rate is zero in the case where per unit time the amount of qNSCs that divides is equal to the amount of qNSCs that are produced from divisions (symmetric + asymmetric divisions).

##### Parameters extraction and evaluation from measured quantities

We define *f_q_, f_a_, f_p_*, and *f_P_* as the fractions of the variables:

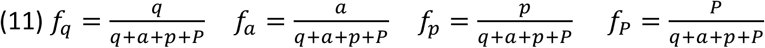

From the fractions in (11) we substitute the variables *q, p* and *P* with *a*, in order to get the following relations:

From equation (7) we get

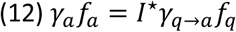

From equations (8) and (9) we get

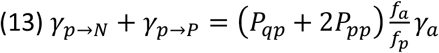

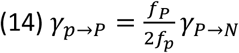

For simplicity we assume that the differentiation rates for aNP and divided aNP are the same. We denote:

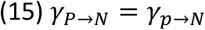

From the relations in (10), (12), (13), (14) and (15), the dynamic parameters are all given by *γ_a_*:

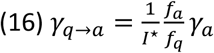

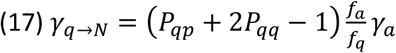

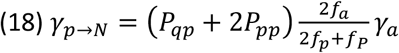

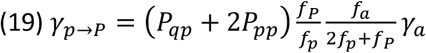

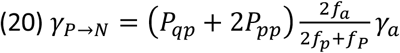

We will next estimate the constants: *γ_a_, P_qq_, P_qp_, P_pp_, f_q_, f_a_, f_p_, f_P_*, and *I*\*.

##### Parameter estimation

###### Estimation of *γ_a_*

From the experimental live tracks (Figure S5) we estimated the duration of the aNSC phase before division. For each fish (Titi, Mimi and Bibi) we counted the number of tracks in which the aNSC phase duration before division was between 0-2, 3-5, 6-8, 9-11, 12-14 and 16 or more days (Table S3, Top). We then estimated the number of tracks that has not divided by a specified time, t (Table S3, Bottom). We then fitted fractions of tracks that has not divided with a decaying exponent of the form *y* = *Ae^−γ_a_t^* (Figure S6B), where *A* is set to 1 (normalized), *t* is the time in days, and *y* is the average fraction of aNSCs tracks that has not divided by time t. Fitting was performed by non-linear least mean square fitting procedure, with 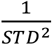 as weights for each data point (at 16 days the weight was set to 10^4^). The best fit for the aNSC decay rate is *γ_a_* = 0.2254 *day*^−1^ with 95% confidence interval of (0.2136, 0.2373).

###### Estimation of probabilities and fractions

From the experimental part (Figure S6A, C and D), we use the following measured values:

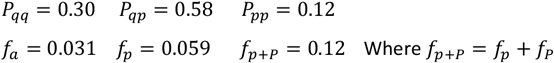

We can then calculate the remaining of the fractions (from the definition in (11)):

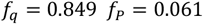

###### Estimation of I*

To simplify the estimation of the mean reduction in qNSC transition rate we assume that if a qNSC is neighbor with one or more aNPs then it is completely inhibited (i.e. it has no chance at all to become aNSC while inhibited). Under this assumption the function *I*(*q, p, P*) gets the form:

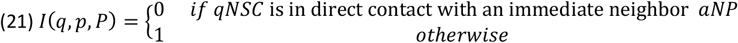

Then, the estimation of *I*\* is the average fraction of qNSC that are not neighboring aNP and therefore are not inhibited. From cell detection of the DM static images from the 4 fish (Figure 1B) we were able to count the fraction of cells that have one or more aNP neighbors (Figure S6F). The fraction of qNSCs that do not neighbor aNP comes out 0.594 ± 0.084 (4 *fish average* ± *STD*). We therefore used the following value *I*\* = 0.59.

From equations (16), (17), (18), (19) and (20), with the estimations and assumptions above, we obtain all the model parameters (Table S4). These parameters are used in a computer simulation as described below.

#### Generation of the NSC lattice model: lattice construction

##### Stochastic simulations on a cell lattice

To test our model, we developed a stochastic simulation platform that simulates the NSC behavior in the DM over 500 days (Video S2). We used a 2D vertex model, similar to the ones defined in (Chiou et al., 2012; Farhadifar et al., 2007; Staple et al., 2010), to generate a dynamic disordered lattice with ~250 cells (Figure 6B). The lattices have periodic boundary conditions to avoid edge effects. The computer simulations are performed on two levels:

1. Tracking cell states and carrying out transitions and events using a probabilistic progression method (Figure S7A -right- and B).
2. Applying changes in morphology according to the events and cell state (Figure S7C-E).

##### Lineage events and LI

The simulations progress with time by calculating the probability for each cell at a certain time step to go through an event in the next time step. The possible cell states and events are as described in the model flowchart (Figures 6A and S7B). The probability for each event, *P_event_*, to take place at the next time step is calculated using the cumulative distribution function (CDF) of an exponential distribution, with λ_*event*_ (the rate for that event) as the distribution rate parameter and *t* = 1 *time step*:

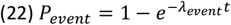

The simulation time interval (i.e. 1 time step) is defined as 1 *day*, and the parameters are used accordingly with the same dimensions (Table S4). Therefore, the probabilities for each event are:

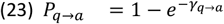

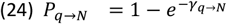

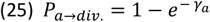

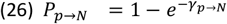

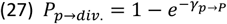

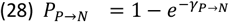

The LI of *qNSC* → *aNSC* by neighboring *aNP* is introduced in the simulation by a modification in the probability of *qNSC* → *aNSC* event (equation (23)) with a term similar to the model function *I*(*q,p,P*), as seen in equations (1) and (2):

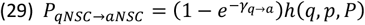

Here *h*(*q, p, P*) is a function of *q, p* and *P* that reduce the rate of *qNSC* transition rate in case of LI. In general, *h* can be a dynamic and continuous function that might depend also on time and space or on other parameters. Consistent with the simplification of *I* (equation (21)) we simplify *h* to be a step function that is 0 if the *qNSC* in question is in direct contact with an immediate neighbor *aNP* (complete inhibition – transition rate is 0 and *P_q→a_* = 0) or 1 otherwise (no inhibition):

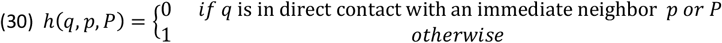

In addition to the model that considers LI, we also consider a model that does not include lateral inhibition. In order to be able to compare between the two models we have to adjust the value of *I*\* to compensate for the overall effect of lack of inhibition. Based on the estimation we set *I*\* = 0.59 for the model with LI, and *I*\* = 1 for the model without LI. This compensation makes sure the fractions of cells in the different states and the percentages following symmetric/asymmetric cell divisions or direct differentiation are the same in both models (Figures 6C and D).

##### Changes in Morphology

There are three main morphology changes in lattice model:

1. Cell division.
2. Cell delamination.
3. Minimization of mechanical energy where each cell state has its own preferred area. Because there are many events of division and delamination at a single time step, a standard 2D vertex model (such as in (Staple et al., 2010)) have a weakness where too many bonds are lost in the process and cells end up having only two bonds (with zero area). The simulation is therefore optimized to reduce the number of bonds lost during cell division and cell delamination (and even sometimes add bonds). The following rules are imposed on the simulations:

##### Cell division

Cell division is done differently for different cell geometries (Figure S7C):

- If the cell consists of 4 bonds then a new bond is added from the middle of the cell’s longest bond to the middle of the opposing bond.
- If the cell consists of 5 bonds then a new bond is add from the middle of the cell’s longest bond to the farthest vertex opposing it.
- If the cell consists of 6 or more bonds then a new bond is added from a random vertex to a vertex which is clockwise far away from it by this formula:

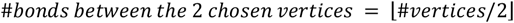

##### Cell delamination

Cell delamination is done in two steps (Figure S7D):

1. If the number of bonds is higher than 4, a T1 transition (intercalation) is applied (Staple et al., 2010).
2. If the number of bonds is 4 the cell is delaminated from the lattice. The delamination process attempts to remove the cell from the lattice with a minimum reduction in the number of bonds from neighboring cells. Furthermore, the algorithm chooses to reduce the number of bonds from the neighbor/s [of the delaminated cell] with the maximum number of bonds. If a cell is surrounded only by cells that each has 4 bonds, the simulation then stops (rarely happens).

##### Mechanical energy minimization

At each time step, as division, delamination and transitions events occur, the lattice relaxes into a new morphological state while minimizing its mechanical energy (Figure S7E). The mechanical energy, *E*, of the 2D lattice at time step t is given by (Farhadifar et al., 2007):

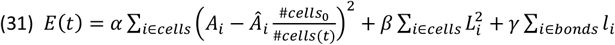

Where *α, β* and *γ* are constant coefficients (see Table S4). The first and second summations over the cells are the contribution to energy given by the area, *A_i_* and perimeter, *L_i_*, respectively. The term *Â_i_* is the preferred area for cell *i*. Note that for each cell state – qNSC, aNSC and aNP, there is a different preferred area *Â_qNSC_, Â_aNSC_* and *Â_aNP_*, respectively (Estimated from experimental data (not shown), Table S4). Our simulations are presented on a fixed size image (Figure 6B), yet the number of cells can vary over time. For that reason the preferred area of cell *i* is scaled by the factor 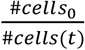, where #*cells*_0_ is the number of cells in the lattice at the initial time step and #*cells*(*t*) is the number of cells in the lattice at time step *t*. In a more realistic model, the division rate is to be regulated by the cell density, keeping the number of cells fixed (with a small deviation). The third summation over the bonds is the energy given by the bond’s tension, associated with its length, *l_i_* (we assume no difference in properties between bonds).

The minimization of equation (31) is done after each cell division and cell delamination and after all transitions in the current time step took place (for example, after all activation events occurred at the time step). Note that the way the energy is minimized is by rearranging all of the vertices in a position which is in the direction that favors the reduction in energy. In our simulation we can control two parameters: the amount of movement in the vertices position (resolution) and the number of iterations in each time the minimization runs. If the resolution is too low (the movement is too large) then the lattice suffers from distortions. If the resolution is high then many iterations are needed in order to make a significance change. Parameters were optimized for highest resolution given the running time constraints. The values of these parameters are available within the code (See Simulation code below).

##### Morphology change due to geometry corrections

We note that our simulation keeps the lattice of cells under the following two restrictions:

1. Any vertex should share 3 cells exactly (no more, no less). If there exist a situation where a vertex is shared more than 3 cells, then a bond is added to the lattice (with a new vertex as well) which makes sure the vertex has only 3 cells sharing it (Figure S7F). The bond is added in the following way: one of the cells that shares the vertex is chosen randomly (with preference to choose an aNSC or aNP). Then a new vertex is added on the connecting axis between the original vertex and the center of the chosen cell. A new bond is then constructed between the original vertex and the new vertex, while the cells that originally shared the original vertex are rearranged with respect to the new created bond.
2. Any cell should have at least 4 bonds. If there exist a situation where a cell has 3 bonds, then the algorithm works out to add a bond to this cell (Figure S7G) without causing a different cell to reduce its number of bonds to 3. Note that the creation of 3 bonds cells are avoided during the simulation.

##### Lattice generation and initial conditions

The cell lattices were generated by starting from an initial 18 by 18 regular hexagonal lattice with periodic boundary conditions. Then, random T1 transitions are performed to break the symmetry of the lattice and small cells are removed from the lattice. For initial conditions we generated a disordered lattice of about 250 cells and set all of the cells to be qNSC, then randomly choose 3.5% of them to be aNSC and different 12% of them to be aNP (where aNPs do not neighbor aNSCs). Afterwards we minimized the lattice mechanical energy in order for the lattice to relax into a morphology that fits better the cells’ state.

##### Simulation code and parameters

We used Matlab software to run the simulations. The code generating the simulations is given in: https://github.com/Udi-Binshtok/NSC_Lattice_model_2020.git

All of the parameters that were used in the simulations are listed in Table S4. These parameters are easily adjustable in the computer simulation code file named defaultparams.m.

## Notes

### Competing Interest Statement

The authors have declared no competing interest.

