## Supplementary figures, tables and discussion for "Dynamic spatiotemporal coordination of neural stem cell fate decisions through local feedback in the adult vertebrate brain"

#### Dray et al., Supplements

Includes

- Supplementary raw data and codes
- 7 Supplementary figure legends
- 4 Supplementary tables
- 2 Supplementary videos
- Supplementary Discussion on Statistics
- Supplementary references

##### Supplementary raw data and codes

Raw data on cell positioning used for the spatial statistics

https://github.com/fcheysson/zebrafish-project

Codes to generate simulations with the NSC lattice model

https://github.com/Udi-Binshtok/NSC_Lattice_model_2020.git

##### Supplementary figures legends

**Figure S1. Distribution of pallial progenitors and proliferation states is similar between hemispheres but differs between anatomical subdivisions, related to Figure 1. (A)** Confocal dorsal view of a whole mount adult telencephalon showing the germinal layer of the pallium in a 3mpf *Tg(gfap:GFP)* fish immunostained for GFP (green, NSCs), PCNA (magenta, proliferating cells) and Sox2 (cyan, NSCs + NPs), and counterstained with DAPI (white). Anterior is to the left. The pallial neuroanatomical subdivisions (Dl: lateral, Dm: medial, Da: anterior pallial domains) are indicated by dotted lines. **(B)** Closeup of the same brain showing qNSCs (Sox2+,Gfap+,PCNA-; green arrows), aNSCs (Sox2+,Gfap+,PCNA+; magenta arrowhead), and aNPs (Sox2+,Gfap-,PCNA+; orange arrowhead). Sox2-,PCNA+ cells (white dotted arrow) are presumably more committed neural progenitors; these cells represent a minority of cell states (Than-Trong et al., 2020). **(C,D)** Quantitative distribution of progenitor states in Dm, **(C)** based on Sox2 expression and **(D)** plotted from four 3 mpf adult fish (1 hemisphere per fish), from an average, per fish, of 1315 qNSCs (s.e.m =170), 82 aNSCs (s.e.m. =11) and 189 aNPs (s.e.m =14) (as in Figure 1D).

**(E-E'')** Semi-automatic cell detection, in the Dm, Da and DI areas, of the stainings for Sox2 and PCNA (color-coded; magenta cells: Gfap-,PCNA+,Sox2+; orange cells: Gfap-,PCNA+,Sox2-). **(E''')** Local density of the dividing cells (PCNA+ and Sox2+,PCNA+ among all Sox2+ cells) in 50µm-diameter spheres, color-coded from 10% (light blue) to 35% (brown) and where the mean is at 20%. Both hemispheres show a very similar density pattern but the three pallial domains are very different (see also (Dray et al., 2015)). Scale bars in A and E are 100µm, in B is 15µm.

**Figure S2. Identification of aNSC doublets as sister cells post-division, and resulting cell detection and filtering method for the spatial analysis of NSC activation events, related to Figure 1. (A-B)** Confocal dorsal view of a whole mount adult telencephalon showing the germinal layer of the pallium in a 3mpf *Tg(gfap:GFP)* fish immunostained for GFP (green), BrdU (red), Sox2 (blue) and PCNA+ZO1 (white, where ZO1 labels apical cell contours), following a 6-hour BrdU pulse and a 18-hour chase. **(A)** Entire views of Dm with focus on BrdU and gfap:GFP; **(B)** closeups from the same brain showing all four markers, with doublets of aNPs (orange arrowheads), doublet of aNSCs (magenta/white arrowheads). **(C)** Identity of BrdU-positive cells after a 6-hour pulse + 18-hour chase. 97% of BrdU+ cells have divided during the chase and are found as doublets (n=137 BrdU+ cell clusters, from 2 fish). Both sister cells are PCNA+ (purple) in 80% of BrdU+ doublets, suggesting that PCNA is detected for some hours after division (n=66 BrdU+ cell clusters from 2 fish). **(D-D'')** Confocal dorsal view of a whole mount adult Dm territory in a 3mpf *Tg(gfap:GFP)* fish immunostained for GFP (green), Sox2 (cyan) and PCNA (magenta) and **(D'-D'')** closeups of the same sample also showing immunostaining with the Zona Occludens marker ZO1. aNSC singlets (magenta arrowhead) are distinguished from doublets (white arrowheads) using the following criteria: proximity of the cell nuclei, similar PCNA staining intensities and similar apical area shape. An aNP singlet is also visible (orange arrowhead). **(E-E'')** Cell detection picture of **(D)** highlighting qNSCs (green dots), aNSCs (magenta dots) and aNPs (orange dots). **(E'-E'')** closeups showing the conversion of a doublet of aNSCs (white arrowheads) into qNSCs. **(F,G)** Cell detection picture of **(D)** showing **(F)** dividing progenitors only (aNSCs and aNPs, magenta and orange spots respectively), and **(G)** aNSCs doublets (white spots) and singlets (magenta spots). **(H)** Resulting schematic representation of the expression of Sox2, Gfap and PCNA along the NSC lineage compared to the BrdU pulse, showing that PCNA staining persists after cytokinesis and can be used to discriminate aNSCs located pre- (singlets) versus post-division (doublets). Scale bars in A is 100µm, in B is 30µm, in D is 50µm and in D'-D'' is 10µm

**Figure S3. Spatial pattern analysis of NSC activation events in 4 independent wildtype brains, related to Figure 1. (A)** Schematics of the principles (top panels) and read-outs (bottom panels) of Besag's L-

function (Besag, 1977) (left), the pair correlation function $g$ (a rescaled derivative of Ripley's $K$-function) (middle), and the $M$-function (right) (Marcon and Puech, 2010; Marcon et al., 2012), assessing correlations between the positions of test cells (red dots) relative to each other within a cell ensemble (green and red dots). $L_{(obs)}(r)$, $g_{(obs)}(r)$ and $M_{(obs)}(r)$ (red lines) are experimental values and, $L_{(H0, mean)}(r)$, $g_{(H0, mean)}(r)$ and $M_{(H0, mean)}(r)$ (black dotted lines) are the means under the null hypothesis Random Labelling (RL), i.e. the state of any cell is independent of other cells and independent of its position (Baddeley et al., 2016). $r$ will be expressed in cell diameters (where 1 is the mean distance between all the cells). Grey regions are the 95% confidence envelopes under the null hypothesis, separating domains of attraction, randomness and repulsion. In short, the $L$- and $M$-functions count the number of test cells within a circled area of radius $r$ around reference cells, while the $g$ function only considers test cells within concentric rings at distance $r$ of the reference cells, which permits to determine the radius of detected interactions. As the $M$-function is adjusted for the frequency of test cells across the entire Dm domain, it helps determine the strength of the interaction. **(B)** Dm surfaces analyzed for each 3mpf adult pallium (4 hemispheres from 4 wildtype fish (WT1 to WT4) (WT4 is the same brain as analyzed in Figures 1E and F), with cell states color-coded. Cell numbers are indicated. **(C)** $L$-functions comparing aNSC activation events with each other at time $t$. $r$ is the mean cell diameter using the mean distances between all cells (qNSC+aNSC+aNP) for each Dm surface. aNSC activation events are randomly spaced relative to each other at time $t$ in all fish. **(D)** $L$-, $g$- and $M$-functions showing that activation events are consistently dispersed relatively to aNPs. Red and dotted black lines are as in (C), green bars indicate the 95<sup>th</sup> centile of the distance to the furthest direct neighbor of aNPs. A $g$-function remaining within the envelope beyond this centile indicates that there is probably no interaction beyond direct neighbors. Reproducibly, NSC activation events are at least two times less likely to occur than predicted by chance within 1-cell diameters from an aNP. **(E)** $L$-, $g$- and $M$-functions showing that aNPs consistently display a clustered pattern at short range (due to the fact that half of them are aNP sister cell doublets, data not shown).

**Figure S4. Spatial pattern analysis of NSC activation events upon Notch blockade, related to Figure 2.** **(A)** Dm surfaces analyzed for each 3mpf adult pallium (4 hemispheres from 4 fish treated with LY for 24h, 3 hemispheres from 3 fish treated with DMSO), with cell states color-coded. Cell numbers are indicated. In each case, the brain on the top row is the same brain as displayed in Figure 2D. **(B)** $L$-functions (Besag, 1977) comparing aNSC activation events with each other at time $t$, where $L_{(obs)}(r)$ (red lines) are experimental values and $L_{(H0, mean)}(r)$ (black dotted lines) are the means under the Random Labelling null hypothesis (CSR). $r$ is the mean cell diameter using the mean distances between all cells (qNSC+aNSC+aNP) for each Dm surface. Grey regions are the 95% confidence envelopes. aNSC activation events are randomly spaced relative to each other at time $t$ in all fish and this is unchanged upon LY treatment. **(C)** $L$-, $g$- and $M$-functions showing that activation events are consistently dispersed relatively to aNPs in DMSO-treated controls, but that this interaction is abolished upon LY treatment. Red and dotted black lines are as in (B), and green bars indicate the 95<sup>th</sup> centile of the distance to the furthest direct neighbor of aNPs. A $g$-function remaining within the envelope beyond this centile indicates that there is probably no interaction beyond direct neighbors. **(D)** $L$-, $g$- and $M$-functions showing that aNPs consistently display a clustered pattern at short range, and that this is unchanged upon LY treatment.

**Figure S5. Track categories (except invariable qNSC tracks) in the three adult fish analyzed, related to Figures 3-5.** **(A)** Dividing and non-dividing track categories and nomenclature, color-coded over time (in days). The “mother cell” (MC in red) is the aNSC at the time point preceding division. **(B)** Representation of all active tracks (ie. either including (a) division(s) event(s) -1st and 2nd columns- or an activation phase -3rd column- or a direct neuronal differentiation -4th column-, in the Dm pallial area of the three fish analyzed (Bibi, Mimi and Titi - further illustrated in Figure 3 and Video S1). Bibi and Mimi and reanalyses of (Than-Trong et al., 2020) incorporating the *mcm5:egfp* staining, and Titi is a new animal. **(C,D)** Quantitative summary of **(B)**. There is a total of 977 qNSC tracks, 102 tracks with 1 division event, 15 with 2 division events, 85 tracks with at least one activated aNSC but no division and 24 tracks losing a NSC through direct differentiation. **(C)** Number of tracks of each state per fish. **(D)** Mean number of NSCs per time point, per fish. **(E)** To focus on aNSCs before division and on first activation events, we filtered the case of consecutive divisions (only the first division event is considered) and converted all aNSCs post-division to a qNSC state. **(E')** Average percentages of NSCs and MCs per time point before and after filtering among all NSCs ; means per time point: “aNSCs before filtering”: 9.6% (s.e.m. 0.6%), “division events”: 1.4% (s.e.m. 0.1%), activation events (“aNSCs before division”): 3.4% (s.e.m. 0.2%), “first division events”: 1.3% (s.e.m. 0.1%).

**Figure S6. Experimental parameters implemented into or used to validate the NSC lattice model, related to Figure 6.** **(A)** Proportion of pre-division activated NSCs at any time point, estimated from the number of aNSC ‘singlets’ (immunostaining experiments, see Figure S2) and from the average number of pre-division aNSCs per time step (live imaging experiments, see Figure S5). From the latter we could also estimate that the average number of dividing NSC per time point (mother cells -MCs-) is 1.3% (s.e.m. 0.1%). **(B)** Estimation of aNSC activation rate ($\gamma_a$). The fraction of remaining aNSCs before division obtained from 88 NSC tracks (blue circles and bars are the average and STD from 3 fish, respectively) (Figure S5 and Table S3) was fitted to a decaying exponential curve (red). The best fit for the activation rate is $\gamma_a = 0.225 \text{ day}^{-1}$ with (0.214 , 0.237) 95% confidence interval. **(C)** Frequency of each division mode, inferred as in (Than-Trong et al., 2020). Symmetric gliogenic divisions produce two NSCs (NSC/NSC), symmetric neurogenic divisions (n/n) produce two aNPs (future neurons, n) that disappear from the germinal sheet, and asymmetric divisions produce one NSC and one aNP (NSC/n). First, to determine the time needed for NSC fates to become apparent, we focused on the most unambiguous fate accessible, ie. the loss of an NSC during the imaging (black arrow). This is due to the loss of the expression of *gfap* (there is almost no cell death nor cell migration (Alunni et al., 2013; Dray et al., 2015; Than-Trong et al., 2020)). **(C')** Cumulative probability distribution of the time between division and 'fate choice' (=loss of *gfap* expression) showing that 90% of neurogenic fates is resolved by 9 days after division for Bibi and 12 days after division for Mimi and Titi. Thus, to estimate division mode we only considered tracks with at least 9 days after division for Bibi and 12 days after division for Mimi and Bibi. **(C'')** Following 53 such tracks, we could estimate that 30.4% (± s.e.m 2.1%) of division events produce two qNSC, 58% (± s.e.m 5.5%) of division events are asymmetric in fate, and 11.9% (± s.e.m 4.3%) of division events have a symmetric neurogenic fate (producing neurons and/or aNPs). **(C''')** Similarly, considering tracks with a direct differentiation happening at least 9 to 12 days after the first time point, we counted 11 direct differentiations. Thus, we could estimate the relative proportions of NSC fates following NSC recruitment (division or direct differentiation) **(C''')** and show that direct differentiations account for 17% (± s.e.m 2.1%) of all events, balancing the gain and loss of NSCs through division (25% (± s.e.m 1.1%) NSC gains via symmetric gliogenic divisions, 9.7% (± s.e.m 5.7%) NSC losses via neurogenic divisions) (similar to Than-Trong et al., 2020). **(D)** Proportion of aNPs among all progenitor cells in Dm, revealed by immunostaining experiments (12.25%, ± s.e.m 1.02%). Among these, about half are present as singlets and half as doublets of aNPs. **(E)** Proportion of aNPs identified as singlets, doublets, triplets or clusters of 4 or more cells (using similar criteria than for the identification of aNSC doublets: similar PCNA staining intensities and similar apical area shape) showing that singlets and doublets of aNPs represent more than 93% of all aNPs. **(F-F'')** Number of neighbors around singlets and doublets of aNPs, and proportion of qNSCs that do not contact an aNP. **(F)** The image shows an example of two aNP singlets (orange, arrow) in contact through edges with 3 qNSCs each (asterisks). Average number of neighbors for singlets and doublets of aNPs (respectively $3.52 \pm$ s.e.m 0.05%, and $4.14 \pm$ s.e.m 0.13%), revealed using immunostainings for ZO1 (yellow) and PCNA (magenta) on four WT fish (Figure S3). **(F''')** Proportion of qNSCs among all qNSCs that are not in contact with any aNPs ($60\% \pm$ s.e.m 5%). Scale bars in F is 10μm.

**Figure S7. Modeling workflow and description of cell fate transition rules in the NSC lattice model, related to Figure 6-7. (A)** Scheme showing the main steps of the model (see methods for details). The two layers of the model include the analytical model (left rectangle) and the NSC lattice model (right rectangle). The analytical model uses as input the experimentally measured values (left inset). The possible transitions in a mean field model are described by the rate equations in the center inset.

Steady state analysis which allows estimation of the transition rates is in the right inset. **(B)** Examples of simulated transitions in two consecutive time steps. **(C)** Schematic of cell division rules. Cell division is carried differently for different cells, depending on the number of bonds. The division rules are designed to leave the daughter cells with at least 4 bonds (see Methods). **(D)** Schematic of a cell delamination process. Cell delamination occurs in two stages: If the cell has 5 or more bonds it reduces its bonds using T1 transition processes (neighbor exchange) until it is left with 4 bonds, only then it delaminates in a way that reduces the minimum amount of bonds in the surrounding cells. The delamination reduces bonds from surrounding cells with 5 or more bonds. Both (C) and (D) are important for proper maintenance of the simulated tissue. **(E)** Examples of cell morphology changes associated with minimization of the lattice mechanical energy, $E$. The mechanical energy depends on 3 terms: cell area (unique area for each cell state, $A_i$), cell perimeter ($L_i$), and bond length ($l_i$). Parameters in the energy term are described in the methods and in Table S4. **(F,G)** Additional morphological corrections required for long term maintenance of the cell lattice. In (E) a new bond is added if a vertex shares more than 3 cells. In (F), a new bond is added to a cell with 3 bonds.

#### Supplementary Videos

**Video S1. 3D animated rendering of the intra-vital imaging, related to Figures 3-5.** The beginning of the movie is a representative animation showing a whole pallial hemisphere imaged intra-vitally in an *casper;Tg(gfap:dTomato);Tg(mcm5:GFP)* double mutant double transgenic 3mpf adult fish (individual fish named Titi) (anterior to the left) followed by a blow-up showing part of the region with cell detection within Dm. The white dots indicate detected cells and the four colored spots (2 magenta and 2 green) highlight a selection of 4 tracks with a division event. Along these tracks, the spot is green when the cell is classified as qNSC and magenta when the cell is classified as aNSC. The second half of the movie shows the 8 successive time points (from day 0 to day 23) and white arrowheads indicate the division events along the 4 selected tracks.

**Video S2. 'NSC lattice model' simulations over 200 time-steps for the model with and without LI, related to Figures 6-7.** One representative simulation out of 18 for each model showing the state changes and morphological changes.

#### Supplementary Tables

**Table S1. Cell states and counts used in the static in vivo analysis, related to Figures 1-2.**

Raw numbers of cells counted per fish (1 hemisphere per animal, within the Dm subregion of the pallium only). Cells were classified as qNSCs (Sox2+, Gfap+), aNSC (Sox2+, Gfap+, PCNA+) or aNP (Sox2+, PCNA+). The last three columns are showing the relative proportions of each cell state within the entire Sox2+ population.

**Table S2. Cell states and counts used in the dynamic in vivo analysis, related to Figures 3-5.**

Raw numbers of the sum of cells counted for the three fish Mimi, Bibi and Titi (1 hemisphere per animal, within the Dm subregion of the pallium only) before (top table) and after (bottom table) filtering (See Figure S5E). The last three columns are showing the relative proportions of each cell state within the qNSC + aNSC population.

**Table S3. Estimation of aNSC decay rate, $\gamma_a$, related to Figure 6 and Figures S6-7.** Top: duration of aNSC tracks. Each row shows the number of tracks dividing within the specified time window, for each fish (col 1-3) and on average (col4) with standard deviation (STD, col 5). Bottom: remaining number of undivided aNSC tracks. Each row is a subtraction of the number of tracks at a specific time, $t$, from the total number of tracks.

**Table S4. NSC lattice model parameters, related to Figures 6-7 and Figure S7.** Summary and values of all the parameters that were used in the simulations.

**Supplementary text (related to STAR Methods -statistics-)**

**Justification of spatial statistics methods**

***The M-function***

While the use of Ripley's $K$-, Besag's $L$- and pair correlation functions is widespread in spatial statistics, the development of the $M$-function is relatively recent, and its use has almost exclusively been circumscribed to economic applications, with only few applications to biology (Fernandez-Gonzalez et al., 2005). However, it presents some improvements to existing measures, mainly that it is a relative, rather than a topographic, measure of spatial concentration. As a consequence, measures of the $M$-function can be interpreted immediately as the relative density of points and give an estimate of the interaction strength exerted by NSCs.

Nevertheless, the $L$- and $g$-functions must not be discarded, as they are complimentary rather than rival to the $M$-function, and they have their advantages for hypothesis testing. First, the $L$-function has approximately stable variance with respect to the range $r$, which makes it a prime candidate for global tests based on simultaneous envelopes, which we used to determine whether an interaction was present. Second, the $g$-function was used to determine the range of the correlation, though other non-cumulative measures (the $Kd$-function (Duranton and Overman, 2005) or the $m$-function (Lang et al., 2020)) could have been used as well. Overall, the measures we used highlight the large statistical toolbox available to the study of spatial patterns in biology.

##### Multiple testing

Care must be taken when interpreting $p$-values resulting from the spatio-temporal analysis of division events (Figure 7). Since testing for each $\Delta t$ time intervals is done on the same samples, the threshold for significance should be adjusted to account for multiple tests and avoid too many false positives. The most common approach is the Bonferroni correction, which adjusts the threshold by dividing it by the number of tests done. Five $\Delta t$ time intervals were considered, so the Bonferroni-adjusted threshold for significance was 0.01 for a desired overall significance level of 0.05. This led, when testing the experimental samples, to the null hypothesis for $\Delta t4$, *i.e.* non-correlation between division events, not being rejected in spite of the low $p$-value (0.026).

However, testing of 18 simulations from the NSC lattice model in the presence of aNP-driven LI reveals that the correlation is statistically significant at $\Delta t4$ time interval, even at this adjusted threshold ($p$-value = 0.009) (Figure 7). Since there was no specific behavior in the model accounting for temporal delay between division events, this behavior must stem from the locally spatial aNP-driven LI. Indeed, the same test applied to 18 simulations in the absence of LI did not recover the interaction at $\Delta t4$ time interval ($p$-value = 0.256) (not shown).

Finally, note the difference between Bonferroni correction and Fisher's method. The former is used in the case of repeated tests on the same samples to avoid too many false positives, while the latter is adapted for replications of the same test on independent samples, to increase the statistical power of the test. While the Bonferroni correction could also be used in the second case to account for repeated tests, it would instead slightly decrease the statistical power (Moran, 2003).

##### Supplementary references

Alunni, A., Krecsmarik, M., Bosco, A., Galant, S., Pan, L., Moens, C.B., Bally-Cuif, L., Ables, J.L.,

- Decarolis, N.A., Johnson, M.A., et al. (2013). Notch3 signaling gates cell cycle entry and limits neural stem cell amplification in the adult pallium. *Development* *140*, 3335–3347.
- Baddeley, A., Rubak, E., and Turner, R. (2016). *Spatial point patterns: methodology and applications with R*. p. 218.
- Besag, J. (1977). Discussion on Dr Ripley's Paper. *J. R. Stat. Soc. Ser. B* *39*, 193–195.
- Dray, N., Bedu, S., Vuillemin, N., Alunni, A., Coolen, M., Krecsmarik, M., Supatto, W., Beaurepaire, E., and Bally-Cuif, L. (2015). Large-scale live imaging of adult neural stem cells in their endogenous niche. *Development* *142*, 3592–3600.
- Duranton, G., and Overman, H.G. (2005). Testing for localization using micro-geographic data. *Rev. Econ. Stud.* *72*, 1077–1106.
- Fernandez-Gonzalez, R., Barcellos-Hoff, M.H., and Ortiz-de-Solórzano, C. (2005). A tool for the quantitative spatial analysis of complex cellular systems. *IEEE Trans. Image Process.* *14*, 1300–1313.
- Lang, G., Marcon, E., and Puech, F. (2020). Distance-based measures of spatial concentration: introducing a relative density function. *Ann. Reg. Sci.* https://doi.org/10.1007/s00168-019-00946-7
- Marcon, E., and Puech, F. (2010). Measures of the geographic concentration of industries: Improving distance-based methods. *J. Econ. Geogr.* *10*, 745–762.
- Marcon, E., Puech, F., and Traissac, S. (2012). Characterizing the relative spatial structure of point patterns. *Int. J. Ecol.* https://doi.org/10.1155/2012/619281
- Moran, M.D. (2003). Arguments for rejecting the sequential bonferroni in ecological studies. *Oikos*.
- Than-Trong, E., Kiani, B., Dray, N., Ortica, S., Simons, B., Rulands, S., Alunni, A., and Bally-Cuif, L. (2020). Lineage hierarchies and stochasticity ensure the long-term maintenance of adult neural stem cells. *Sci. Adv.* *6*, eaaz5424.

Figure S1

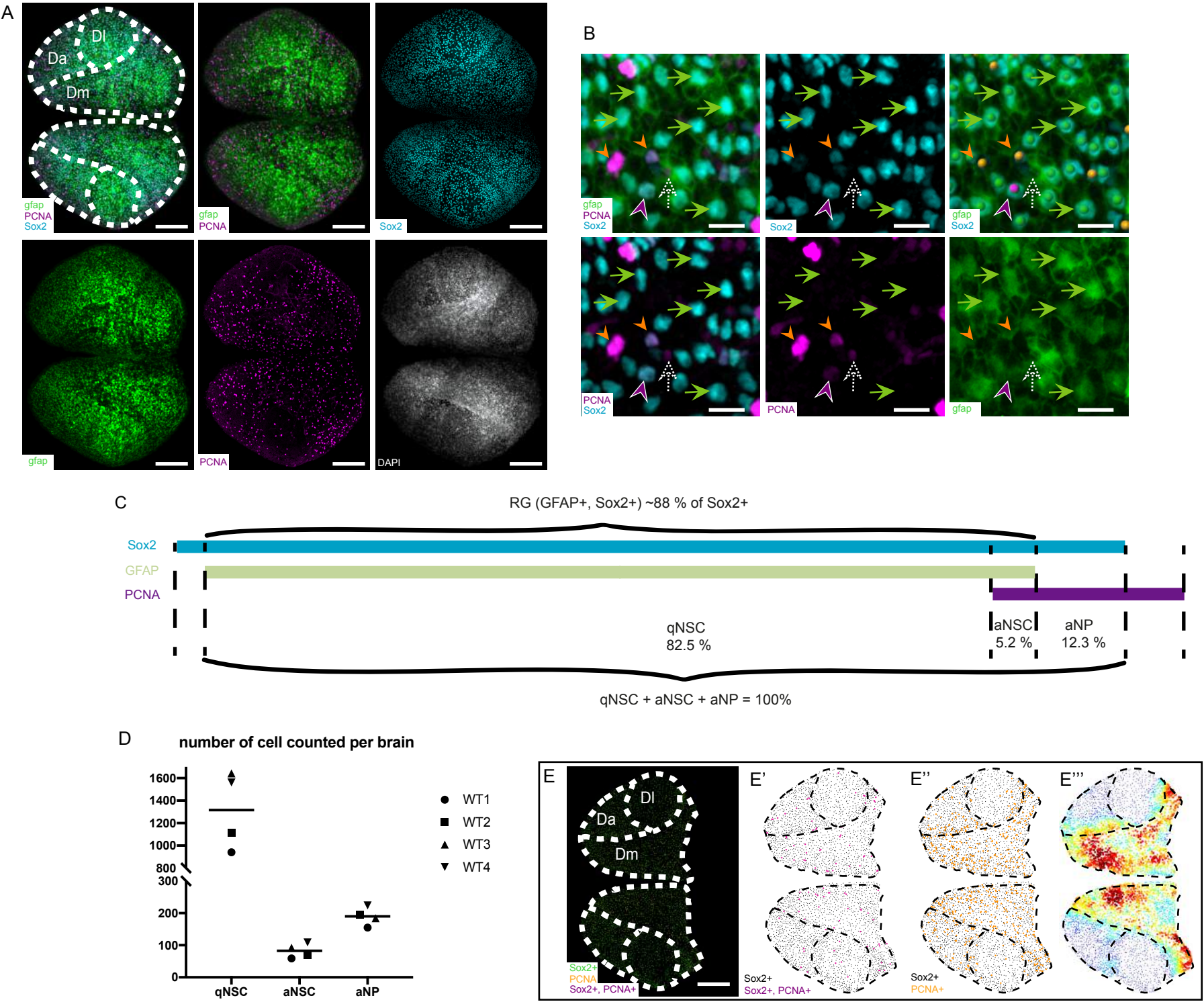

### Figure S2

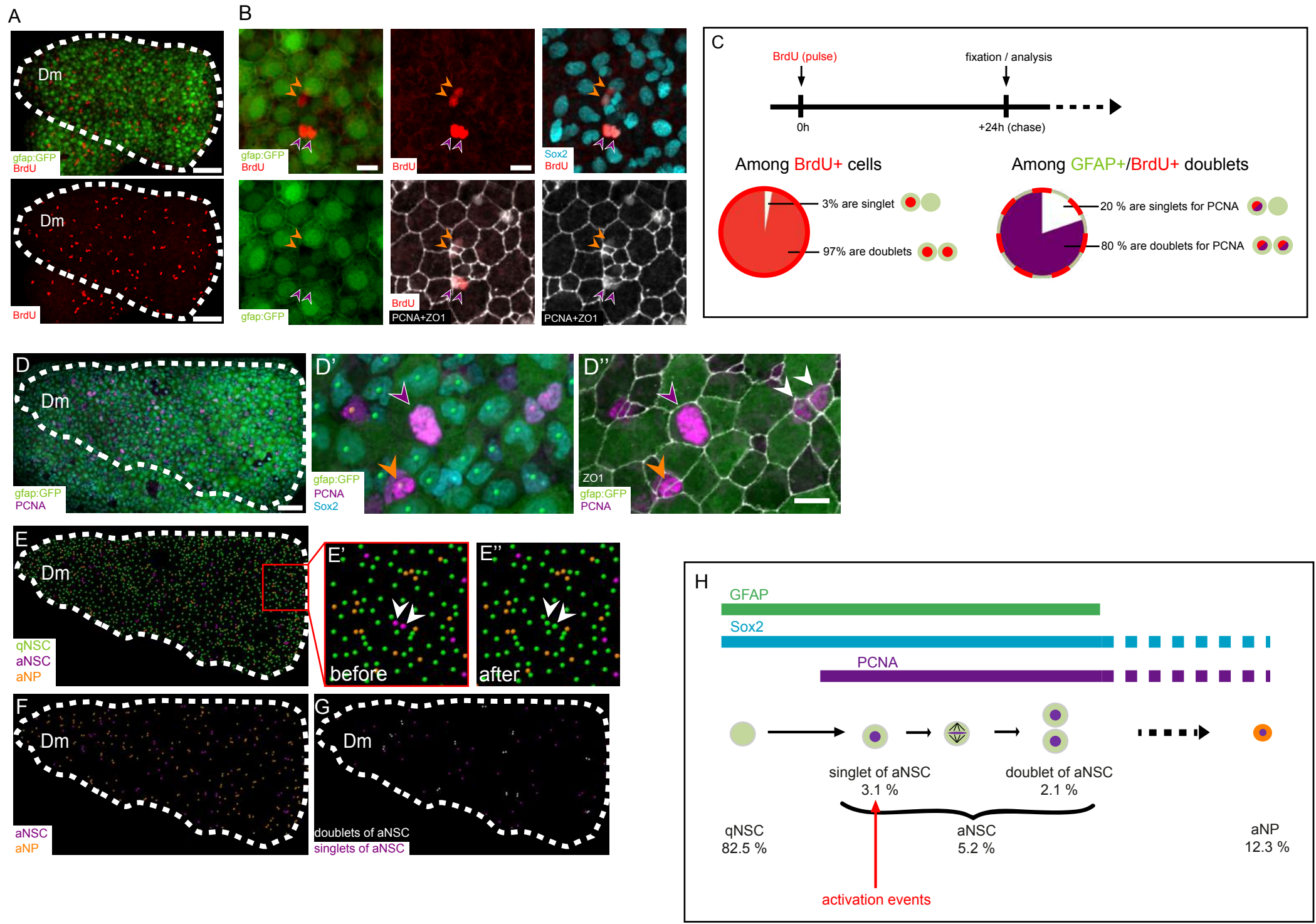

Figure S3

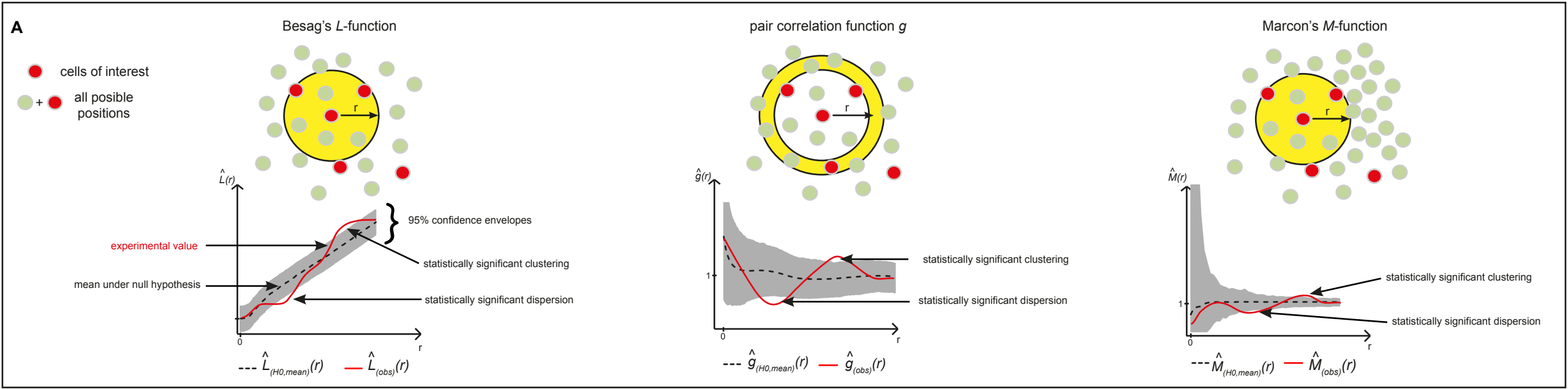

**B**

● qNSC
● aNSC
● aNP

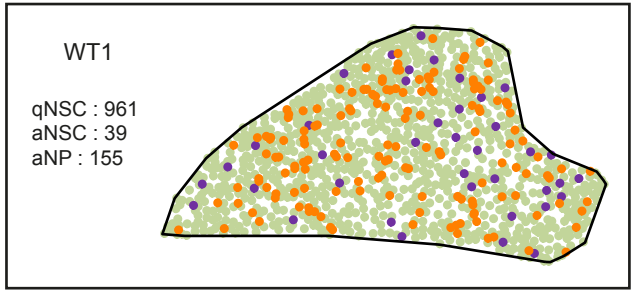

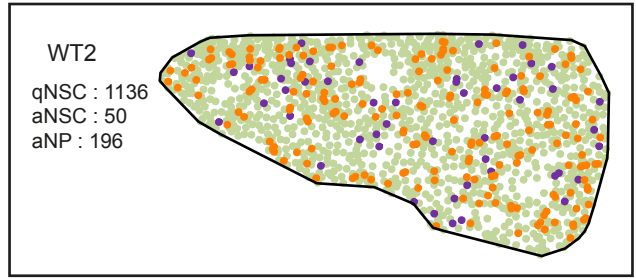

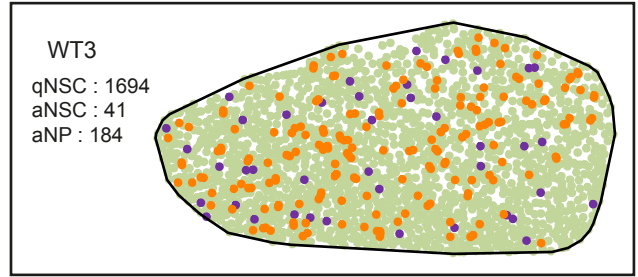

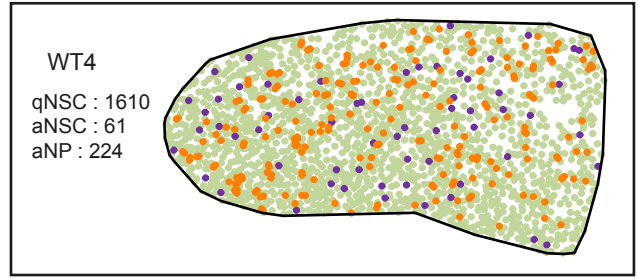

**C**

aNSC vs aNSC

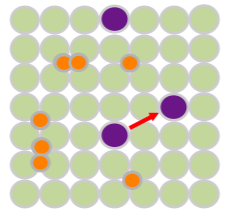

--- $\hat{L}_{(H0,mean)}(r)$ --- $\hat{L}_{(obs)}(r)$

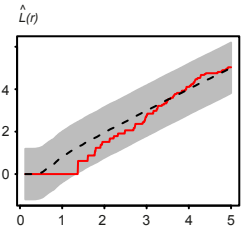

**D**

aNSC vs aNP

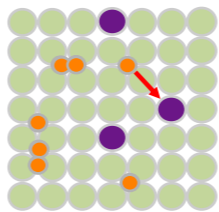

--- $\hat{L}_{(H0,mean)}(r)$ --- $\hat{L}_{(obs)}(r)$

--- $\hat{g}_{(H0,mean)}(r)$ --- $\hat{g}_{(obs)}(r)$

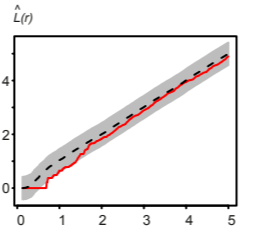

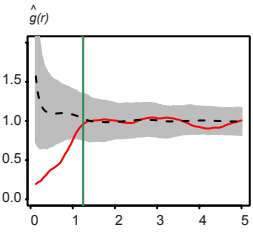

**E**

aNP vs aNP

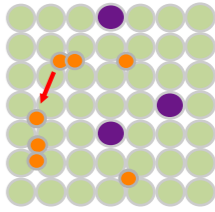

--- $\hat{L}_{(H0,mean)}(r)$ --- $\hat{L}_{(obs)}(r)$

--- $\hat{g}_{(H0,mean)}(r)$ --- $\hat{g}_{(obs)}(r)$

--- $\hat{M}_{(H0,mean)}(r)$ --- $\hat{M}_{(obs)}(r)$

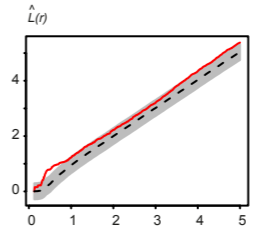

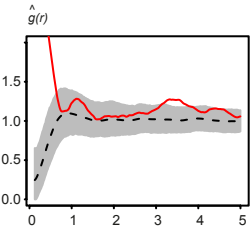

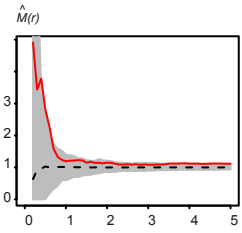

$r$ (cell diameter)

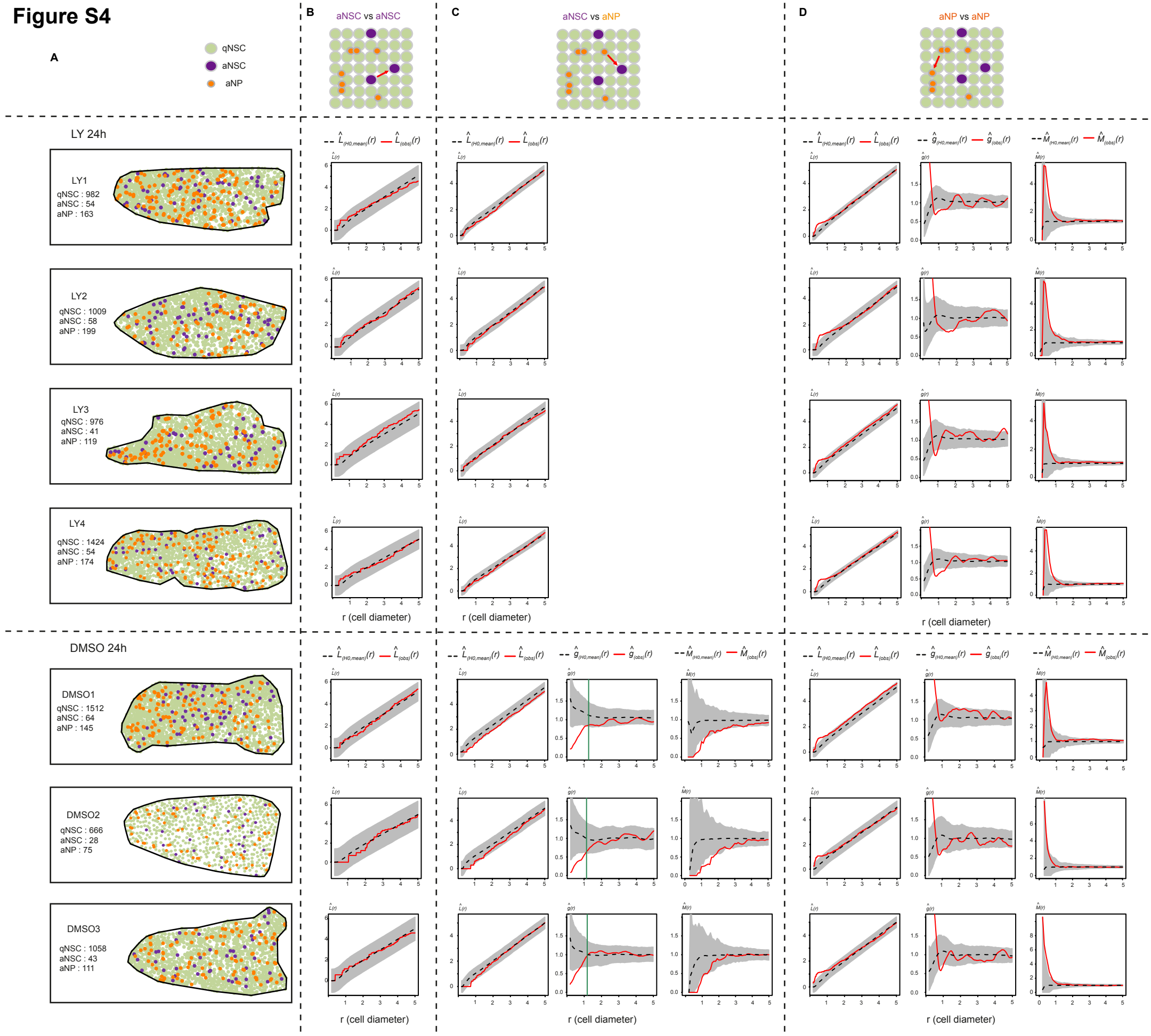

**Figure S5**

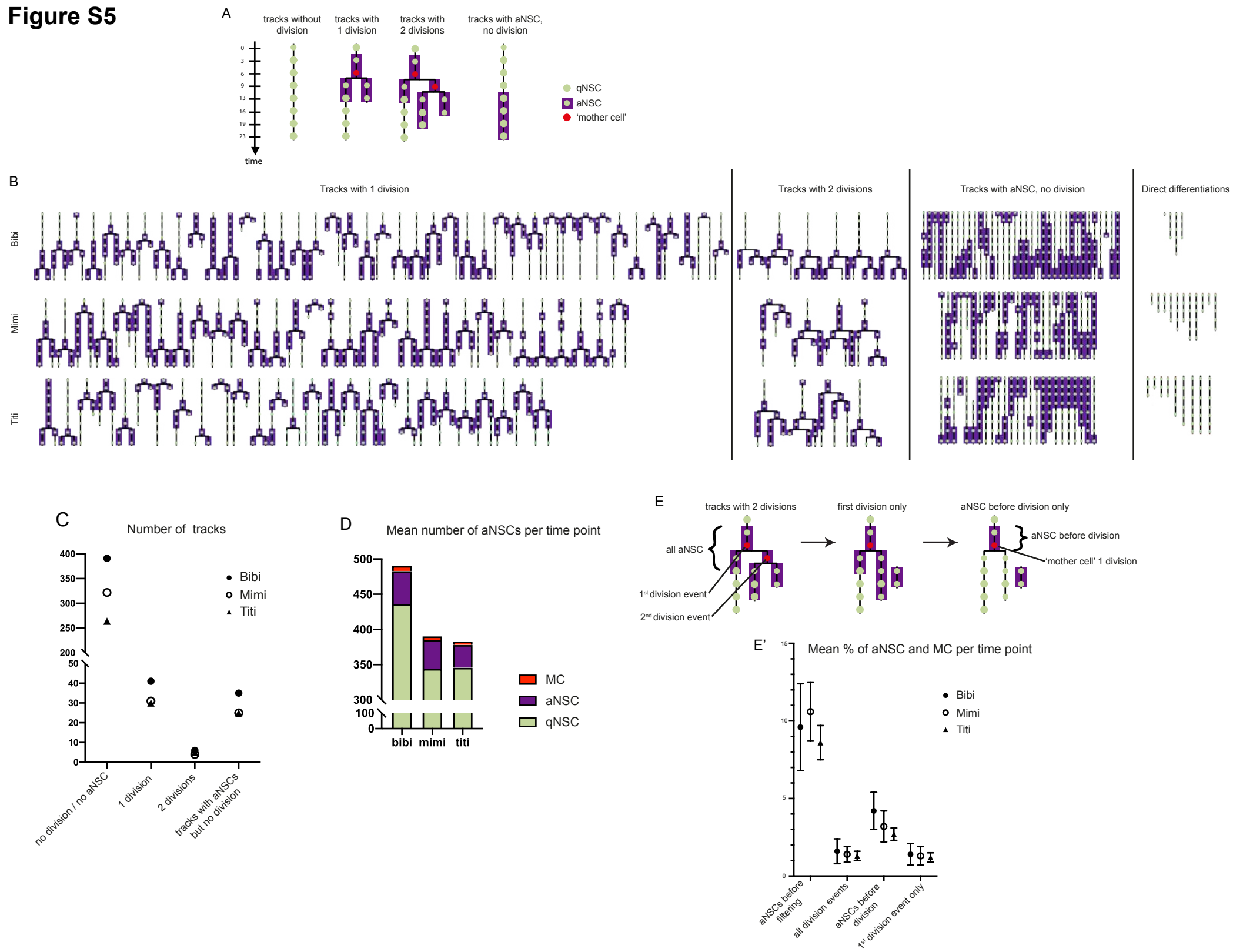

Figure S6

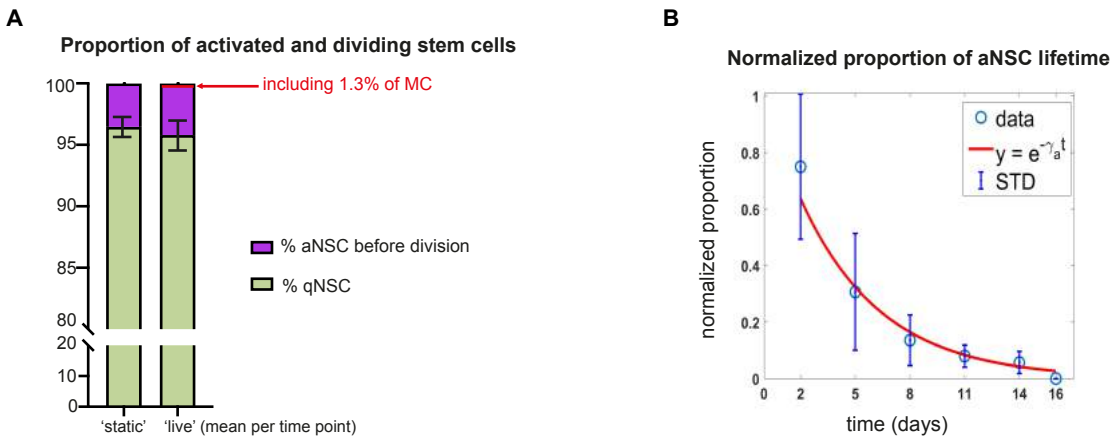

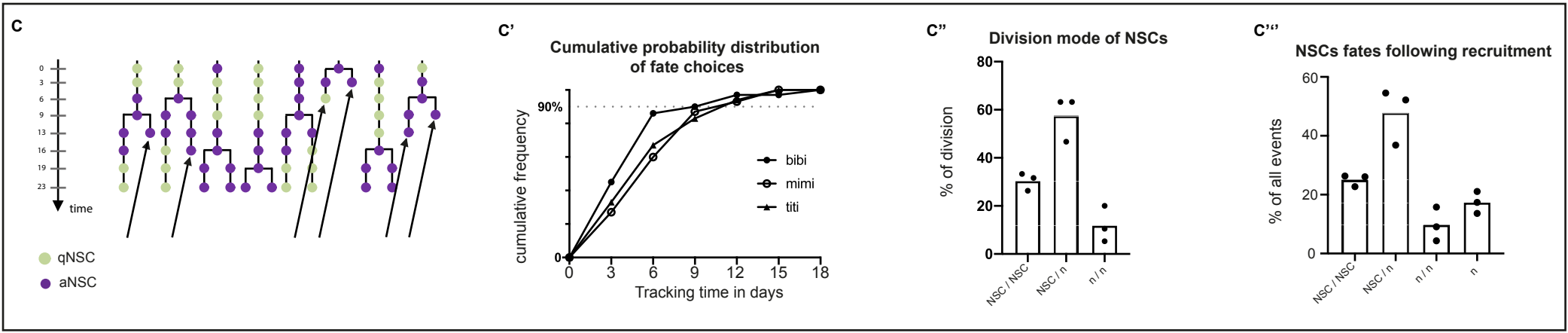

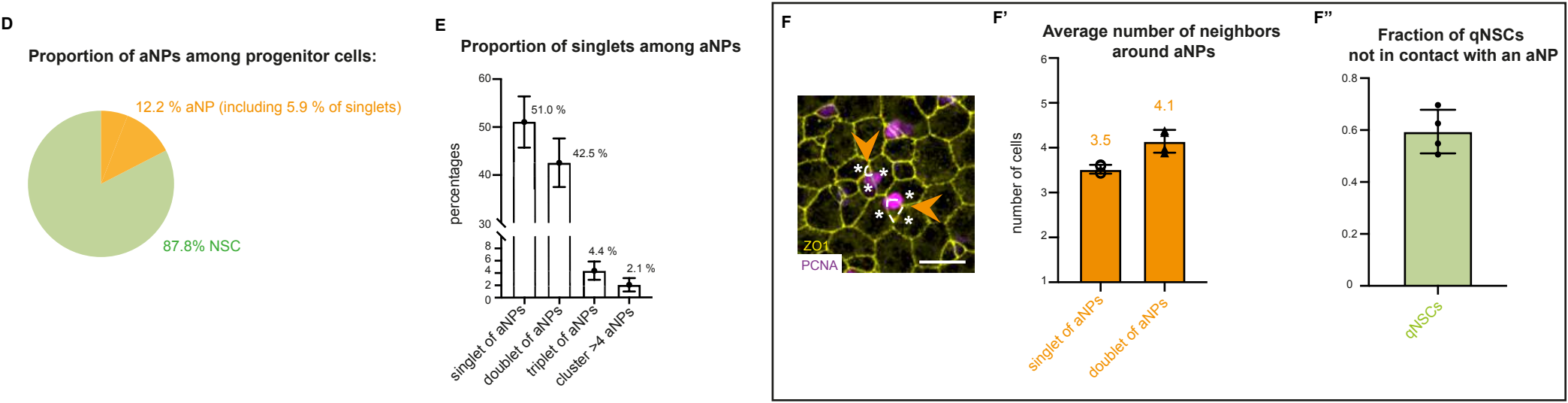

**Figure S7**

**A**

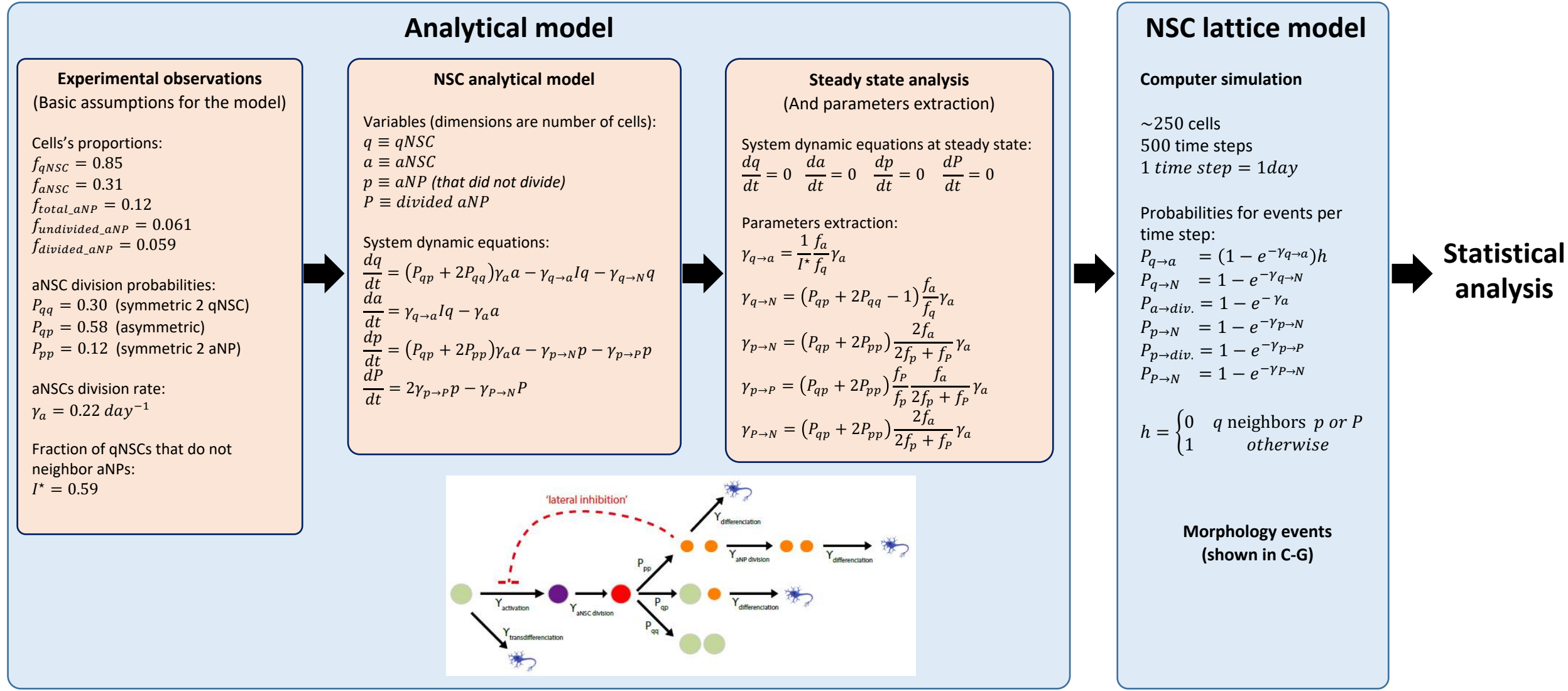

**B**

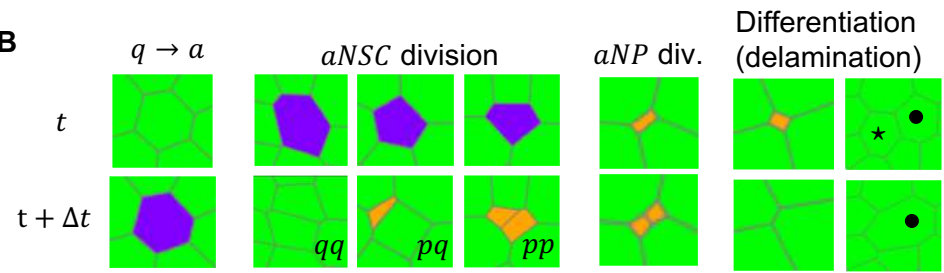

**C**

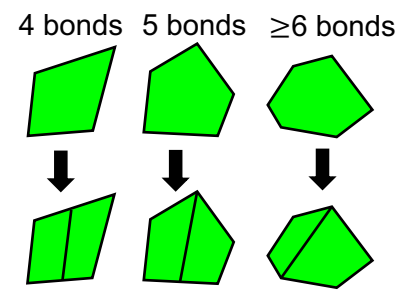

**D**

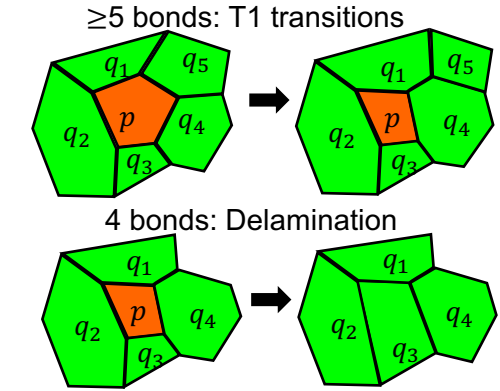

**E**

$$E = \alpha \sum_{i \in \text{cells}} (A_i - \hat{A}_i)^2 + \beta \sum_{i \in \text{cells}} L_i^2 + \gamma \sum_{i \in \text{bonds}} l_i$$

Minimizing cell's perimeter and bonds' length

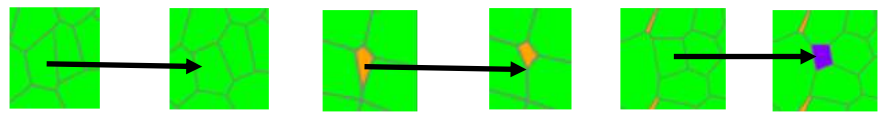

**F**

$\geq 4$ cells shares a vertex:
add bond (and vertex)

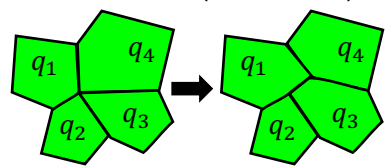

**G**

3 bonds: add bond

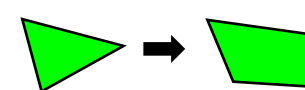

**Table S1.** Cell types and counts used in the static in vivo analysis, related to Figures 1-2.

| all aNSCs | among Sox2+ cells only | # qNSC | # aNSC | # aNP | total<br>number of<br>cells | % among: qNSC+aNSC+aNP |  |  |
| --- | --- | --- | --- | --- | --- | --- | --- | --- |
|  |  |  |  |  |  | % aNSC | % aNP | % qNSC |
| WT | WT1 | 941.0 | 59.0 | 155.0 | 1155.0 | 5.1 | 13.4 | 81.5 |
|  | WT2 | 1116.0 | 70.0 | 196.0 | 1382.0 | 5.1 | 14.2 | 80.8 |
|  | WT3 | 1643.0 | 92.0 | 184.0 | 1919.0 | 4.8 | 9.6 | 85.6 |
|  | WT4 | 1562.0 | 109.0 | 224.0 | 1895.0 | 5.8 | 11.8 | 82.4 |
|  | average WT | **1315.5** | **82.5** | **189.8** | **1587.8** | **5.2** | **12.3** | **82.6** |
|  | std | **340.6** | **22.4** | **28.6** | **380.2** | **0.4** | **2.0** | **2.1** |
|  | s.e.m | **170.3** | **11.2** | **14.3** | **190.1** | **0.2** | **1.0** | **1.1** |
| ly24h | LY1 | 938.0 | 98.0 | 163.0 | 1199.0 | 8.2 | 13.6 | 78.2 |
|  | LY2 | 977.0 | 90.0 | 99.0 | 1166.0 | 7.7 | 8.5 | 83.8 |
|  | LY3 | 946.0 | 71.0 | 119.0 | 1136.0 | 6.3 | 10.5 | 83.3 |
|  | LY4 | 1344.0 | 134.0 | 174.0 | 1652.0 | 8.1 | 10.5 | 81.4 |
|  | average WT | **1051.3** | **98.3** | **138.8** | **1288.3** | **7.6** | **10.8** | **81.7** |
|  | std | **195.9** | **26.4** | **35.6** | **243.9** | **0.9** | **2.1** | **2.5** |
|  | s.e.m | **97.9** | **13.2** | **17.8** | **121.9** | **0.4** | **1.1** | **1.3** |
| DMSO24h | DMSO1 | 1484.0 | 92.0 | 145.0 | 1721.0 | 5.3 | 8.4 | 86.2 |
|  | DMSO2 | 644.0 | 50.0 | 75.0 | 769.0 | 6.5 | 9.8 | 83.7 |
|  | DMSO3 | 1016.0 | 85.0 | 111.0 | 1212.0 | 7.0 | 9.2 | 83.8 |
|  | average WT | **1048.0** | **75.7** | **110.3** | **1234.0** | **6.3** | **9.1** | **84.6** |
|  | std | **420.9** | **22.5** | **35.0** | **476.4** | **0.9** | **0.7** | **1.4** |
|  | s.e.m | **243.0** | **13.0** | **20.2** | **275.0** | **0.5** | **0.4** | **0.8** |

| singlet of<br>aNSCs | among Sox2+ cells only | # qNSC | # aNSC | # aNP | total<br>number of<br>cells | % among: qNSC+aNSC+aNP |  |  |
| --- | --- | --- | --- | --- | --- | --- | --- | --- |
|  |  |  |  |  |  | % aNSC | % aNP | % qNSC |
| WT | WT1 | 961.0 | 39.0 | 155.0 | 1155.0 | 3.4 | 13.4 | 83.2 |
|  | WT2 | 1136.0 | 50.0 | 196.0 | 1382.0 | 3.6 | 14.2 | 82.2 |
|  | WT3 | 1694.0 | 41.0 | 184.0 | 1919.0 | 2.1 | 9.6 | 88.3 |
|  | WT4 | 1610.0 | 61.0 | 224.0 | 1895.0 | 3.2 | 11.8 | 85.0 |
|  | average WT | **1350.3** | **47.8** | **189.8** | **1587.8** | **3.1** | **12.3** | **84.7** |
|  | std | **357.3** | **10.0** | **28.6** | **380.2** | **0.7** | **2.0** | **2.7** |
|  | s.e.m | **178.7** | **5.0** | **14.3** | **190.1** | **0.3** | **1.0** | **1.3** |
| ly24h | LY1 | 982.0 | 54.0 | 163.0 | 1199.0 | 4.5 | 13.6 | 81.9 |
|  | LY2 | 1009.0 | 58.0 | 99.0 | 1166.0 | 5.0 | 8.5 | 86.5 |
|  | LY3 | 976.0 | 41.0 | 119.0 | 1136.0 | 3.6 | 10.5 | 85.9 |
|  | LY4 | 1424.0 | 54.0 | 174.0 | 1652.0 | 3.3 | 10.5 | 86.2 |
|  | average WT | **1097.8** | **51.8** | **138.8** | **1288.3** | **4.1** | **10.8** | **85.1** |
|  | std | **218.0** | **7.4** | **35.6** | **243.9** | **0.8** | **2.1** | **2.2** |
|  | s.e.m | **109.0** | **3.7** | **17.8** | **121.9** | **0.4** | **1.1** | **1.1** |
| DMSO24h | DMSO1 | 1512.0 | 64.0 | 145.0 | 1721.0 | 3.7 | 8.4 | 87.9 |
|  | DMSO2 | 666.0 | 28.0 | 75.0 | 769.0 | 3.6 | 9.8 | 86.6 |
|  | DMSO3 | 1058.0 | 43.0 | 111.0 | 1212.0 | 3.5 | 9.2 | 87.3 |
|  | average WT | **1078.7** | **45.0** | **110.3** | **1234.0** | **3.6** | **9.1** | **87.3** |
|  | std | **423.4** | **18.1** | **35.0** | **476.4** | **0.1** | **0.7** | **0.6** |
|  | s.e.m | **244.4** | **10.4** | **20.2** | **275.0** | **0.0** | **0.4** | **0.4** |

**Table S2.** Cell types and counts used in the dynamic in vivo analysis, related to Figures 3-5.

| before filtering (see Figure S5E) |  |  |  |  |  |  |
| --- | --- | --- | --- | --- | --- | --- |
| time point | # qNSC | # aNSC | # MC | % qNSC | % aNSC | % MC |
| t1 | 1,131.0 | 89.0 | 17.0 | 92.7 | 7.3 | 1.4 |
| t2 | 1,133.0 | 99.0 | 12.0 | 92.0 | 8.0 | 1.0 |
| t3 | 1,131.0 | 106.0 | 16.0 | 91.4 | 8.6 | 1.3 |
| t4 | 1,131.0 | 115.0 | 16.0 | 90.8 | 9.2 | 1.3 |
| t5 | 1,129.0 | 120.0 | 17.0 | 90.4 | 9.6 | 1.4 |
| t6 | 1,112.0 | 142.0 | 27.0 | 88.7 | 11.3 | 2.2 |
| t7 | 1,113.0 | 153.0 | 21.0 | 87.9 | 12.1 | 1.7 |
| t8 | n/a | n/a | n/a | n/a | n/a | n/a |
| sum all t | 7,880.0 | 824.0 | 126.0 |  |  |  |
| mean per t | 1,125.7 | 117.7 | 18.0 | 90.6 | 9.4 | 1.4 |
| std | 9.1 | 22.9 | 4.8 | 1.7 | 1.7 | 0.4 |
| s.e.m | 3.4 | 8.7 | 1.8 | 0.7 | 0.7 | 0.1 |

| after filtering (see Figure S5E) |  |  |  |  |  |  |
| --- | --- | --- | --- | --- | --- | --- |
| time point | # qNSC | # aNSC | # MC | % qNSC | % aNSC | % MC |
| t1 | 1,183.0 | 37.0 | 17.0 | 97.0 | 3.0 | 1.4 |
| t2 | 1,199.0 | 33.0 | 11.0 | 97.3 | 2.7 | 0.9 |
| t3 | 1,199.0 | 38.0 | 15.0 | 96.9 | 3.1 | 1.2 |
| t4 | 1,209.0 | 37.0 | 16.0 | 97.0 | 3.0 | 1.3 |
| t5 | 1,204.0 | 45.0 | 14.0 | 96.4 | 3.6 | 1.1 |
| t6 | 1,197.0 | 57.0 | 24.0 | 95.5 | 4.5 | 1.9 |
| t7 | 1,216.0 | 50.0 | 15.0 | 96.1 | 3.9 | 1.2 |
| t8 | n/a | n/a | n/a | n/a | n/a | n/a |
| sum all t | 8,407.0 | 297.0 | 112.0 |  |  |  |
| mean per t | 1,201.0 | 42.4 | 16.0 | 96.6 | 3.4 | 1.3 |
| std | 10.4 | 8.6 | 4.0 | 0.7 | 0.7 | 0.3 |
| s.e.m | 3.9 | 3.3 | 1.5 | 0.2 | 0.2 | 0.1 |

**Table S3. Estimation of aNSC decay rate, $\gamma_a$**

| # of aNSC tracks dividing between: | titi | mimi | Bibi | average | STD |
| --- | --- | --- | --- | --- | --- |
| 0 to 2 days | 9 | 6 | 7 | 7.3 | 1.5 |
| 3 to 5 days | 12 | 10 | 17 | 13 | 3.6 |
| 6 to 8 days | 1 | 7 | 7 | 5 | 3.5 |
| 9 to 11 days | 0 | 3 | 2 | 1.7 | 1.5 |
| 12 to 14 days | 0 | 2 | 0 | 0.67 | 1.2 |
| 16 days or more | 1 | 1 | 3 | 1.7 | 1.2 |
| **Total** | **23** | **29** | **36** | **29.3** |  |
| ***Remaining aNSC tracks by time t***<br>**= (total # tracks)**<br>**– (# of tracks divided by time t)** |  |  |  |  |  |
| t = 2 days | 14 | 23 | 29 | 22 | 7.5 |
| t = 5 days | 2 | 13 | 12 | 9 | 6.1 |
| t = 8 days | 1 | 6 | 5 | 4 | 2.6 |
| t = 11 days | 1 | 3 | 3 | 2.3 | 1.2 |
| t = 14 days | 1 | 1 | 3 | 1.7 | 1.2 |
| t = 16 days | 0 | 0 | 0 | 0 | 0 |

**Table S4. NSC lattice model parameters.**

| Parameter | Description | value |
| --- | --- | --- |
| $f_q$ | Fraction of qNSCs (from the total cells) | 0.849 |
| $f_a$ | Fraction of aNSCs (from the total cells) | 0.031 |
| $f_p$ | Fraction of aNPs (from the total cells) | 0.059 |
| $f_P$ | Fraction of divided aNPs (from the total cells) | 0.061 |
| $I^*$ | Fraction of qNSCs that are not in contact with any aNP (from the total qNSCs). Represent the fraction of qNSCs that are not inhibited by aNPs. | 0.59 with LI<br>1 without LI |
| $\gamma_{q \rightarrow a}$ | qNSC activation rate ($day^{-1}$) | 0.014 with LI<br>0.0080 without LI |
| $\gamma_{q \rightarrow N}$ | qNSC differentiation rate ($day^{-1}$) | 0.0014 |
| $\gamma_a$ | aNSC division rate ($day^{-1}$) | 0.22 |
| $\gamma_{p \rightarrow N}$ | aNP differentiation rate ($day^{-1}$) | 0.062 |
| $\gamma_{p \rightarrow P}$ | aNP division rate ($day^{-1}$) | 0.032 |
| $\gamma_{P \rightarrow N}$ | aNP differentiation rate ($day^{-1}$) | 0.062 |
| $P_{qq}$ | probability for symmetric division giving 2 qNSC | 0.30 |
| $P_{qp}$ | probability for asymmetric division giving 1 qNSC 1 aNP | 0.58 |
| $P_{pp}$ | probability for symmetric division giving 2 aNP | 0.12 |
| $\alpha$ | Energy minimization area factor in equation (31) | 40 |
| $\beta$ | Energy minimization perimeter factor in equation (31) | 2.5 |
| $\gamma$ | Energy minimization bonds factor in equation (31) | 5 |
| $\hat{A}_{qNSC}$ | Energy minimization qNSC preferred area. $\hat{A}$ in (31) | $\frac{\text{initial lattice area}}{\text{initial num. of cells}}$ |
| $\hat{A}_{aNSC}$ | Energy minimization aNSC preferred area. $\hat{A}$ in (31) | $0.35\hat{A}_{qNSC}$ |
| $\hat{A}_{aNP}$ | Energy minimization mother cell preferred area. $\hat{A}$ in (31) | $0.10\hat{A}_{qNSC}$ |
